# Spatial protein quality control requires a functional GET pathway to prevent proteotoxicity

**DOI:** 10.64898/2026.10.15.759444

**Authors:** Chloe A. Langridge, Emily M. Sontag

## Abstract

The proper management of misfolded proteins is an essential biological process that is required for the survival of all organisms. Previous work has demonstrated that misfolded proteins can be spatially sequestered into membrane-less compartments to facilitate their degradation. Soluble cytoplasmic misfolded proteins are sequestered in the juxtanuclear quality control compartment (JUNQ), and insoluble cytoplasmic misfolded proteins are sequestered into the insoluble protein deposit (IPOD). It is still unknown if there are upstream targeting pathways necessary for regulating JUNQ and IPOD formation. Here we investigated the role of ER targeting on JUNQ and IPOD formation. We find that the GET pathway is necessary for JUNQ formation. Inefficiencies in the GET pathway contribute to premature IPOD formation, and a defective GET pathway inhibits JUNQ formation altogether. Notably, we find the GET pathway’s role in JUNQ formation is required for cell survival and loss of GET pathway function is only toxic under proteotoxic stress. Overall, we identify a new role for the GET pathway in regulating spatial PQC and demonstrate that spatial quality control is essential for cell health and survival.

## Introduction

Maintaining correctly folded proteins is an essential biological process for cell and organismal health. Under cellular stress, when proteins are most susceptible to misfolding, molecular chaperones are produced and sort misfolded proteins into protein quality control compartments. These membrane-less compartments sequester misfolded proteins to facilitate degradation^1–3^ In the cytosol, misfolded proteins are sequestered into small cytosolic compartments along the cortical ER called Q-bodies^4^. Over time, Q-bodies move along the ER and coalesce into larger inclusions at the perinuclear ER. Eventually Q-bodies coalesce into a large inclusion at the nuclear-vacuolar junction called the juxtanuclear quality control (JUNQ) compartment^1,2,4,5^. A peripheral inclusion called the insoluble protein deposit (IPOD) also forms to sequester insoluble or amyloid-like misfolded proteins^3,5–8^. It remains unknown if the formation of Q-bodies or the JUNQ contribute to cell survival.

Q-body localization and formation is dependent on the J-domain chaperone proteins Ydj1 and Hlj1^2,4^, both located at the ER^9–12^. Farnesylation of Ydj1 anchors it to the cytosolic surface of the ER membrane^10,11^. Hlj1 is a tail anchored protein with a C-terminal hydrophobic transmembrane domain^9^ that is inserted into the ER membrane by the ER-targeting SND and GET pathways^13,14^. While the chaperones required to induce Q-body formation and maturation are known, the upstream pathways that contribute to Q-body regulation remain unclear. Specifically, the role of ER targeting pathways in the regulation of Q-bodies has yet to be characterized.

While SND and GET pathways for ER targeting are highly overlapping, there are key differences in their substrates^13,14^. The GET pathway preferentially binds a single, highly hydrophobic, transmembrane domain in the C-terminal “tail” of tail-anchored proteins^15–18^. Tail-anchored proteins are inserted in a stepwise handoff by the members of the GET pathway^19,20^. The function of the GET pathway is not essential under unstressed conditions but has been proposed to be essential under stress conditions^21–23^.

More recently, various chaperones of the GET pathway have been shown to have moonlighting functions in protein quality control. Under oxidative stress, Get3 oligomerizes by cysteine oxidation and functions as a holdase chaperone^24,25^. During the induction of stress granules, Get5 is imported into the nucleus and interacts with stress granule marker Pab1^26^. Sgt2 is interacts with amyloid proteins^27^ and is necessary for the recruitment of other GET chaperones and heat shock proteins to GET bodies under glucose starvation^28^.

In this study, we investigated the role of the GET pathway on inclusion formation utilizing EGFP-VHL as a constitutively misfolded reporter protein. Previous studies have proposed Sgt2 and other members of the GET pathway may be important for proteostasis because they have roles outside of tail anchored protein insertion^23,27^. We find that under proteotoxic stress the GET pathway is essential for Q-body and JUNQ formation. When we break the GET pathway at any stage in the tail anchored protein insertion process, Q-body formation is inhibited, and the JUNQ does not form. Additionally, we find loss of GET pathway proteins is toxic but only under proteotoxic stress when JUNQ formation is lost. This indicates that JUNQ formation is required for cell survival under proteotoxic stress, and the GET pathway plays a crucial role in regulating JUNQ formation.

## Results

### Sgt2 prevents VHL but not NES-LuciTs inclusion formation at 30°C

Sgt2 interacts with other GET pathway chaperones in its N-terminus (GET), heat shock proteins in its tetracopeptide repeat (TPR) domain, and tail anchored proteins in its C-terminus (Sti1) (Fig. 1a). Previous work has identified Sgt2 as an interactor of amyloid substrates in its Sti1 domain and proposed Sgt2 may have a key role in PQC^27^. We utilized well known cytoplasmic spatial PQC substrates to determine if deletion of *SGT2* influenced PQC pathways of soluble substrates or if these interaction reports were specific to amyloid substrates.

**Figure 1:**
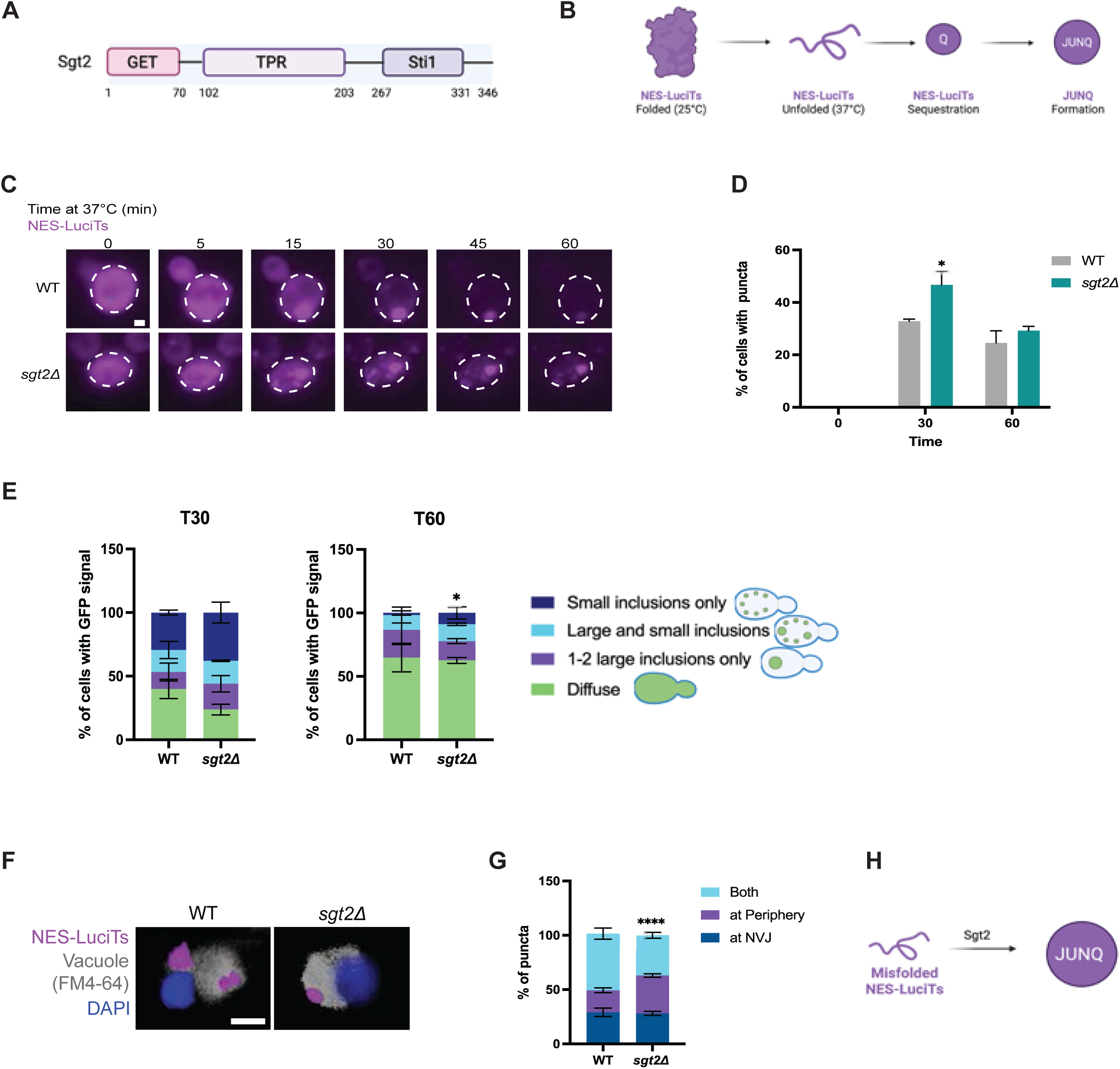
Sgt2 sorts misfolded NES-LuciTs to the JUNQ. Schematics showing the domains of the Sgt2 protein in (A) and a temperature-sensitive mutant of Luciferase with a nuclear export signal tag (NES-LuciTs) that misfolds when temperature shifts to 37°C in (B). (C) Time-resolved live cell microscopy of wildtype BY4741 and *SGT2* deletion strains expressing NES-LuciTs at 37°C (magenta). Representative still frames at the times shown. Three biological replicates were performed. (D) Percentage of cells with puncta over time. Fisher’s test: at T30 *p-value=0.0372, at T60 p-value=0.3361. (E) Categorization of cells expressing NES-LuciTs at 37°C after 30 minutes and 60 minutes of timelapse. Fisher’s test: at T30 p-value=0.1248, at T60 *p-value=0.0369. (F) Representative images of FM4-64 vacuole staining of live cells (gray) with Hoechst 33342 staining (blue) after 30 minutes heat shock. (G) Quantification of inclusion location from (F). Inclusions in proximity to the nuclear-vacuolar junction were categorized at NVJ. Inclusions away from nuclear-vacuolar junction on either the nuclear or vacuolar side were categorized as at periphery. Cells with inclusions at least one inclusion near the nuclear-vacuolar junction and at least one inclusion away from the nuclear-vacuolar junction were categorized as both. Fisher’s test: ****p-value=<0.0001. Yeast growth of wildtype BY4741 and *SGT2* deletion strains in 2% galactose and 2% raffinose media at 30C (H) and 37C (I). Area under the curve analysis of (H, I) in (J, K) respectively. For (I) one-way ANOVA: WT vs. WT + NES-LuciTs ##p-value=0.0017, *sgt2Δ* vs. *sgt2Δ* + NES-LuciTs ###p-value=0.0002. All error bars represent standard error of the mean (SEM). (L) schematic showing Sgt2 sorts misfolded NES-LuciTs to the JUNQ.

While *sgt2Δ* did not strongly impact inclusion formation of NES-LuciTs (Fig. 1c-e, S1b), NES-LuciTs steady state levels (Fig. S1a) or growth (Fig. 1h-k, S1c-h), we did observe changes in localization of the inclusions. In *sgt2Δ* cells, NES-LuciTs more often forms only peripheral inclusions rather than simultaneously forming both peripheral and nuclear-vacuolar junction (NVJ) localized inclusions like wild type cells (Fig. 1f-g, l) suggesting Sgt2 aids in targeting NES-LuciTs to the JUNQ (Fig. 1h). We next wanted to determine if Sgt2 only targets NES-LuciTs or if Sgt2 plays a broader role in targeting misfolded proteins to the JUNQ. We utilized EGFP-VHL to track the role of Sgt2 in the sequestration of terminally misfolded proteins (Fig. 2a) Interestingly, loss of Sgt2 induces VHL inclusion formation prior to heat shock (Fig. 2b-d, S2b) without altering VHL steady state levels (Fig. S2a). Prior to and during heat shock, VHL preferentially forms 1-2 large inclusions in *sgt2Δ* cells rather than gradually transitioning from several small inclusions to 1-2 large inclusions over the course of an hour like wild type cells (Fig. 2d). This indicates that Sgt2 could be involved in the initial formation of Q-bodies as the small, dynamic puncta fail to form in the *sgt2Δ* cells. The inclusions that form prior to heat shock in *sgt2Δ* cells equally localize to the nuclear vacuolar junction (NVJ) or periphery but rarely form in both places at once (Fig. 2e-f). This data suggests that loss of Sgt2 induces VHL sequestration into PQC compartments without heat shock (Fig. 2g). Despite the formation of these inclusions, we did not observe major changes in yeast growth compared to wild type cells (Fig. S2c-h), showing that the inclusions formed are not toxic or protective. Taken together, these data indicate that Sgt2 can distinguish between thermally unstable proteins like temperature-sensitive luciferase, and constitutively misfolded proteins like VHL.

**Figure 2:**
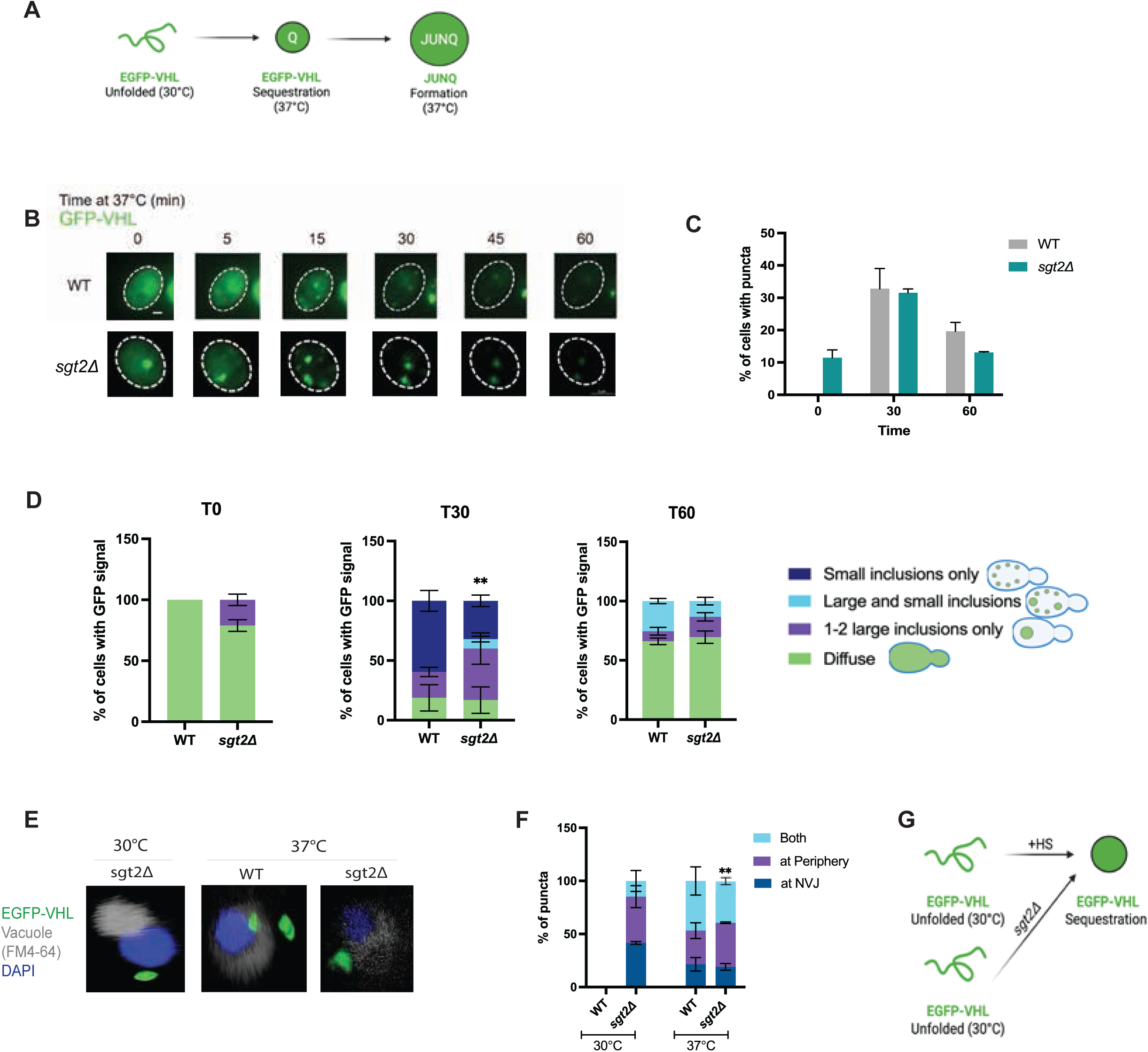
Loss of *SGT2* induces EGFP-VHL sequestration without heat shock. (A) schematic showing terminally misfolded EGFP-VHL and its sequestration pathway at 37°C. (B) Time-resolved live cell microscopy of wildtype and *SGT2* deletion strains expressing EGFP-VHL at 37°C (green). Representative still frames at the times shown. Three biological replicates were performed. (C) Percentage of cells with puncta over time. Fisher’s test: p-value>0.05. (D) Categorization of inclusions in cells expressing EGFP-VHL at 37°C after 30 minutes and 60 minutes of timelapse. Fisher’s test and Bonferroni correction: at T30 **p-value=0.005. (E) Representative images of FM4-64 vacuole staining of live cells (gray) with Hoechst 3342 staining (blue) prior to and after 30 minutes of heat shock. (F) Quantification of inclusion localization from (G). Inclusions in proximity to the nuclear-vacuolar junction were categorized at NVJ. Inclusions away from nuclear-vacuolar junction on either the nuclear or vacuolar side were categorized as at periphery. Cells with inclusions at least one inclusion near the nuclear-vacuolar junction and at least one inclusion away from the nuclear-vacuolar junction were categorized as both. Fisher’s test and Bonferroni correction: *p-value=0.0448. Yeast growth of wildtype BY4741 and *SGT2* deletion strains in 2% galactose and 2% raffinose media at 30°C (G) and 37°C (I). Area under the curve analysis of (G) in (H) and (I) in (J). In (H), one-way ANOVA: WT vs. WT + EGFP-VHL ###p-value=0.0005, *sgt2Δ* vs. *sgt2Δ* + EGFP-VHL #p-value=0.0160, In (J), one-way ANOVA: WT vs. WT + EGFP-VHL ###p-value=0.006, *sgt2Δ* vs. *sgt2Δ* + EGFP-VHL #p-value=0.0186.

### Substrates of the GET pathway co-aggregate with VHL in the IPOD in the absence of Sgt2

We next wanted to determine what could be causing the formation of VHL inclusions in *sgt2Δ* cells. Because we observed VHL inclusion formation prior to heat shock, we predicted deleting *SGT2* may prematurely induce the heat shock response. However, we did not observe changes in Hsp70 levels (Fig. S2a).

Several studies have demonstrated tail anchored proteins can aggregate in the cytosol if they are not properly inserted into the ER membrane. Notably, the tail-anchored protein Sed5 utilizes the GET pathway and Sgt2 for its membrane insertion^20^. Because Sed5 mislocalizes to the cytoplasm instead of the ER in *sgt2Δ* cells, we wanted to determine if Sed5 was still mislocalized under proteotoxic stress conditions, and if the mislocalized Sed5 is aggregated in the cytosol (Fig. 3a).

**Figure 3:**
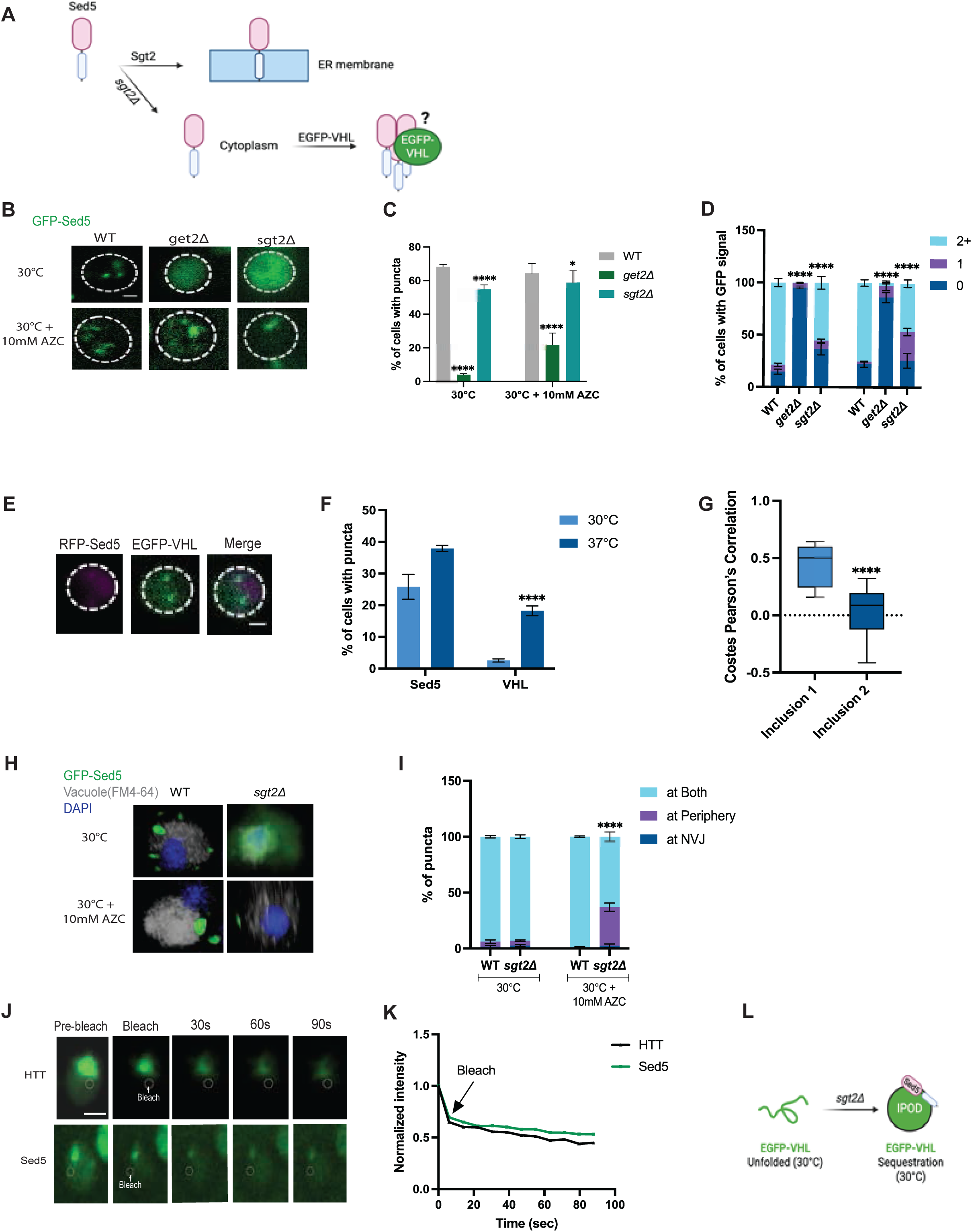
Mislocalized Sed5 co-aggregates with VHL to form an IPOD. (A) schematic showing Sgt2 aids in the insertion of Sed5 into the ER membrane. (B) Live cell microscopy of wildtype, *GET2* deletion, and *SGT2* deletion strains expressing GFP-Sed5 +/- 1 hour of 10mM AZC treatment. Representative images are shown. Three biological replicates were performed. (C) Percentage of cells with puncta over time. Fisher’s test and Bonferroni correction: ****p-value<0.0001. (D) Categorization of number of inclusions per cell in cells expressing GFP-Sed5. Fisher’s test and Bonferroni correction: ****p-value<0.0001. (E) Live cell microscopy of *SGT2* deletion cells expressing EGFP-VHL and mCherry-Sed5. Representative images are shown. Three biological replicates were performed. (F) Percentage of cells from (E) with puncta before and after 30 minutes of heat shock. Fisher’s test: VHL 30°C vs. VHL 37°C ****p-value<0.0001. (G) Costes Pearson’s correlation co-efficient of inclusions in cells with 2 EGFP-VHL inclusions. Welch’s t-test: ****p-value<0.0001. (H) Representative images of FM4-64 vacuole staining of live cells (gray) with Hoechst 3342 staining (blue) +/- 1 hour 10mM AZC exposure. (I) Quantification of inclusion localization from (H). Inclusions in proximity to the nuclear-vacuolar junction were categorized at NVJ. Inclusions away from nuclear-vacuolar junction on either the nuclear or vacuolar side were categorized as at periphery. Cells with inclusions at least one inclusion near the nuclear-vacuolar junction and at least one inclusion away from the nuclear-vacuolar junction were categorized as both. Fisher’s test: ****p-value<0.0001. (J) Representative images of wildtype cells expressing exon1 expanded huntingtin or *sgt2Δ* cells expressing GFP-Sed5. Photobleached region of cytosol indicated with white circle. Fluorescence of cells was measured for 90 seconds following bleach. (K) Normalized intensity of aggregate before and after bleaching adjacent cytosol. Aggregate intensity was normalized to intensity of aggregate before bleaching. If the fluorescent intensity of background >1, aggregate intensity was also normalized to background. Mean of three biological replicates of at least 10 cells from each replicate was plotted. (L) schematic showing in the absence of *SGT2*, Sed5 co-aggregates with misfolded proteins under proteotoxic stress triggering premature IPOD formation.

We expressed GFP-Sed5 in wild type, *sgt2Δ,* and *get2Δ* cells with and without the presence of 10mM L-Azetidine-2-carboxylic acid (AZC), a proline analog that induces protein misfolding, to trigger proteotoxic stress. Because of Sed5’s role in vesicle fusion, it forms several puncta in wild type cells that indicate vesicle formation and fusion not misfolded protein accumulation. Under unstressed conditions, GFP-Sed5 forms less puncta and is more diffuse in *sgt2Δ* cells albeit not as dramatically as in *get2Δ* cells (Fig. 3b-d, S3c) in line with previous reports^31^. We also observed a decrease in Sed5 steady state levels as expected (Fig. S3a). After AZC treatment, we observed a decrease in cells with diffuse Sed5 in *sgt2Δ* and *get2Δ* cells compared to wild type cells treated with AZC. In fact, we saw an increase in the number of cells forming one punctum (Fig. 3b-d). We predicted the single Sed5 punctum could be the cause of EGFP-VHL inclusion formation and that was triggering premature inclusion formation.

To determine if Sed5 inclusions colocalize with PQC compartments in *sgt2Δ* cells, we co-expressed mCherry-Sed5 and EGFP-VHL in the same cells. At both 30°C and 37°C we observed high variation in colocalization between Sed5 and VHL (Fig. S3b). However, we noticed at 37°C when EGFP-VHL formed multiple inclusions, RFP-Sed5 would often colocalize with one VHL inclusion but not the other. We also noticed this colocalization often occurred with a VHL inclusion at the cell periphery. We split the puncta of cells forming at least two EGFP-VHL puncta into two groups. One group does not have co-localization of Sed5 and VHL as seen by a lower Pearson’s correlation coefficient of that inclusion and one that does have colocalization shown by a higher Pearson’s correlation coefficient of that inclusion. We found a significant difference between the two groups (Fig. 3e-g) suggesting Sed5 shows preference for one PQC compartment over the other.

We next wanted to determine if the Sed5 inclusion preferentially localized to the JUNQ or the IPOD. In wild type cells, Sed5 forms several inclusions localized both at the periphery and at the NVJ with and without AZC treatment. However, in *sgt2Δ* cells treated with AZC, we found a dramatic increase in the number of Sed5 puncta localized exclusively at the periphery suggesting Sed5 localizes to the IPOD in *sgt2Δ* cells (Fig. 3h-i). To confirm the peripheral Sed5 punctum was an IPOD, we photobleached the cytosol of cells with one Sed5 punctum. We measured the fluorescent intensity of the punctum for 90 seconds to monitor cytosolic exchange with the punctum. Like our HTT IPOD control, the Sed5 punctum remains static confirming its IPOD-like nature (Fig. 3j-k, S3d). Together, this data suggests the mislocalized clients of Sgt2 can co-aggregate with misfolded proteins causing premature IPOD formation (Fig. 3l).

### Blocking the GET binding domain of Sgt2 inhibits misfolded protein inclusion formation

Previous work has demonstrated overexpression of *SGT2* can interfere with the efficiency of the GET pathway. We therefore wanted to determine if increasing steady state levels of Sgt2 thereby altering the stoichiometry of the GET pathway also alters VHL inclusion formation (Fig. 4a), or if only *sgt2Δ* has this effect. To test this, we co-expressed EGFP-VHL and *SGT2*-DsRed from a centromeric plasmid in *sgt2Δ* cells to induce *SGT2* overexpression. We observed inclusion formation prior to heat shock but notably less often than in *sgt2Δ* cells (Fig. 4b-d, S4a). These inclusions localize to either the NVJ or periphery but rarely both, like *sgt2Δ* cells (Fig. 4e-f). Like *sgt2Δ* cells, when we overexpress *SGT2*, inclusion formation after heat shock favors 1-2 large inclusions rather than transitioning from several small inclusions to 1-2 large inclusions like in wild type cells (Fig. 4d). We did not observe major changes in growth or steady state levels of VHL compared to wild type cells (Fig. 4g-j, S2a, S4b-c). This data suggests that increasing or decreasing Sgt2 protein levels alters VHL inclusion formation (Fig. 4k).

**Figure 4:**
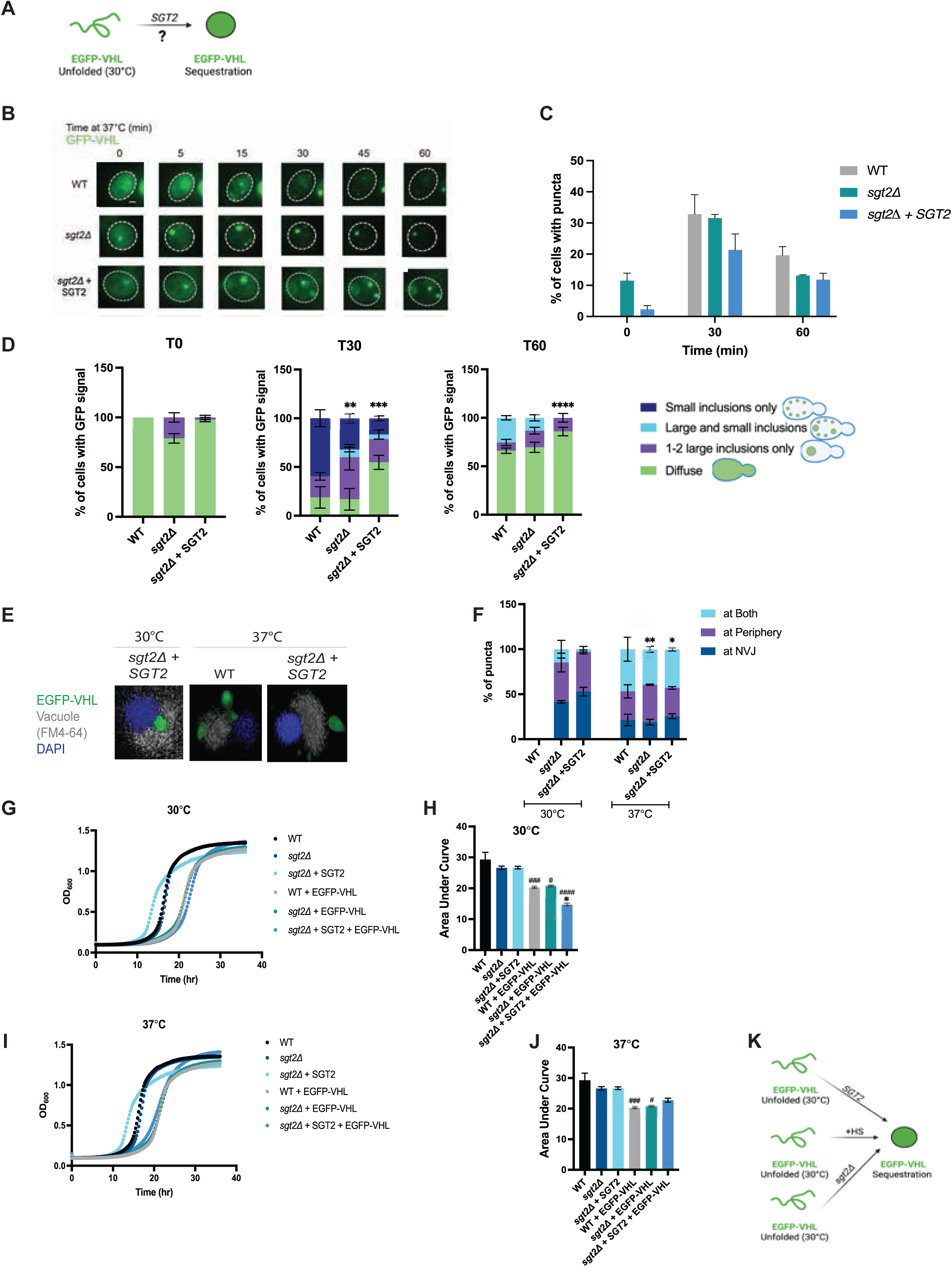
*SGT2* overexpression reduces premature IPOD formation. (A) schematic showing the effect of *SGT2* overexpression on VHL sequestration is unknown. (B) Time-resolved live cell microscopy of wildtype and *SGT2* deletion and overexpression strains expressing EGFP-VHL at 37°C (green). Representative still frames at the times shown. Three biological replicates were performed. (C) Percentage of cells with puncta over time. Fisher’s test: p-value>0.05. (D) Categorization of inclusions in cells expressing EGFP-VHL at 37°C after 30 minutes and 60 minutes of timelapse. Fisher’s test and Bonferroni correction: at T30 ***p-value=0.0003, at T60 ****p-value<0.0001. (E) Representative images of FM4-64 vacuole staining of live cells (gray) with Hoechst 3342 staining (blue) prior to and after 30 minutes of heat shock. (F) Quantification of inclusion localization from (G). Inclusions in proximity to the nuclear-vacuolar junction were categorized at NVJ. Inclusions away from nuclear-vacuolar junction on either the nuclear or vacuolar side were categorized as at periphery. Cells with inclusions at least one inclusion near the nuclear-vacuolar junction and at least one inclusion away from the nuclear-vacuolar junction were categorized as both. Fisher’s test and Bonferroni correction: p-value<0.05. Yeast growth of wildtype BY4741 and *SGT2* deletion, and *SGT2* overexpression strains in 2% galactose and 2% raffinose media at 30°C (G) and 37°C (I). Area under the curve analysis of (G) in (H) and (I) in (J). In (H), one-way ANOVA: WT vs. WT + EGFP-VHL ###p-value=0.0005, *sgt2Δ* vs. *sgt2Δ* + EGFP-VHL #p-value=0.0160, *sgt2Δ* + *SGT2* vs. *sgt2Δ* + *SGT2* + EGFP-VHL ####p-value>0.0001, WT + EGFP-VHL vs. *sgt2Δ* + *SGT2* + EGFP-VHL *p-value=0.0193. In (J), one-way ANOVA: WT vs. WT + EGFP-VHL ###p-value=0.006, *sgt2Δ* vs. *sgt2Δ* + EGFP-VHL #p-value=0.0186. (K) schematic showing EGFP-VHL is sequestered when *SGT2* is lost or supplemented above wild type levels.

We hypothesized the overexpression of *SGT2* may be causing non-productive binding of VHL to Sgt2’s Sti1 domain and contributing to increased VHL aggregation. To test this, we mutated each domain of *SGT2* individually (Fig. 5a) and co-expressed the mutants with EGFP-VHL. We used point mutations previously identified in the literature to reduce binding of Get5 (C39S)^32^, heat shock proteins (R171A/R175A)^19,33,34^, and tail anchored proteins (I286D)^35^. Surprisingly, mutating the GET binding domain (C39S) rather than the Sti1 domain greatly reduced EGFP-VHL inclusion formation at 30°C and 37°C (Fig. 5c, S5a). Additionally, the C39S GET binding domain mutant reduces growth at 30°C and 37°C but only when VHL is present (Fig. 5e-j, Fig. S5b-e). Mutating any of the three domains altered the localization of the inclusions at 37°C (Fig. 5b-dd). Together, these data indicate that a functional GET binding domain is necessary for Q-body formation (Fig. 5i) Further, this is the first indication that the inability of misfolded proteins to form PQC compartments is detrimental to cell health and survival.

**Figure 5:**
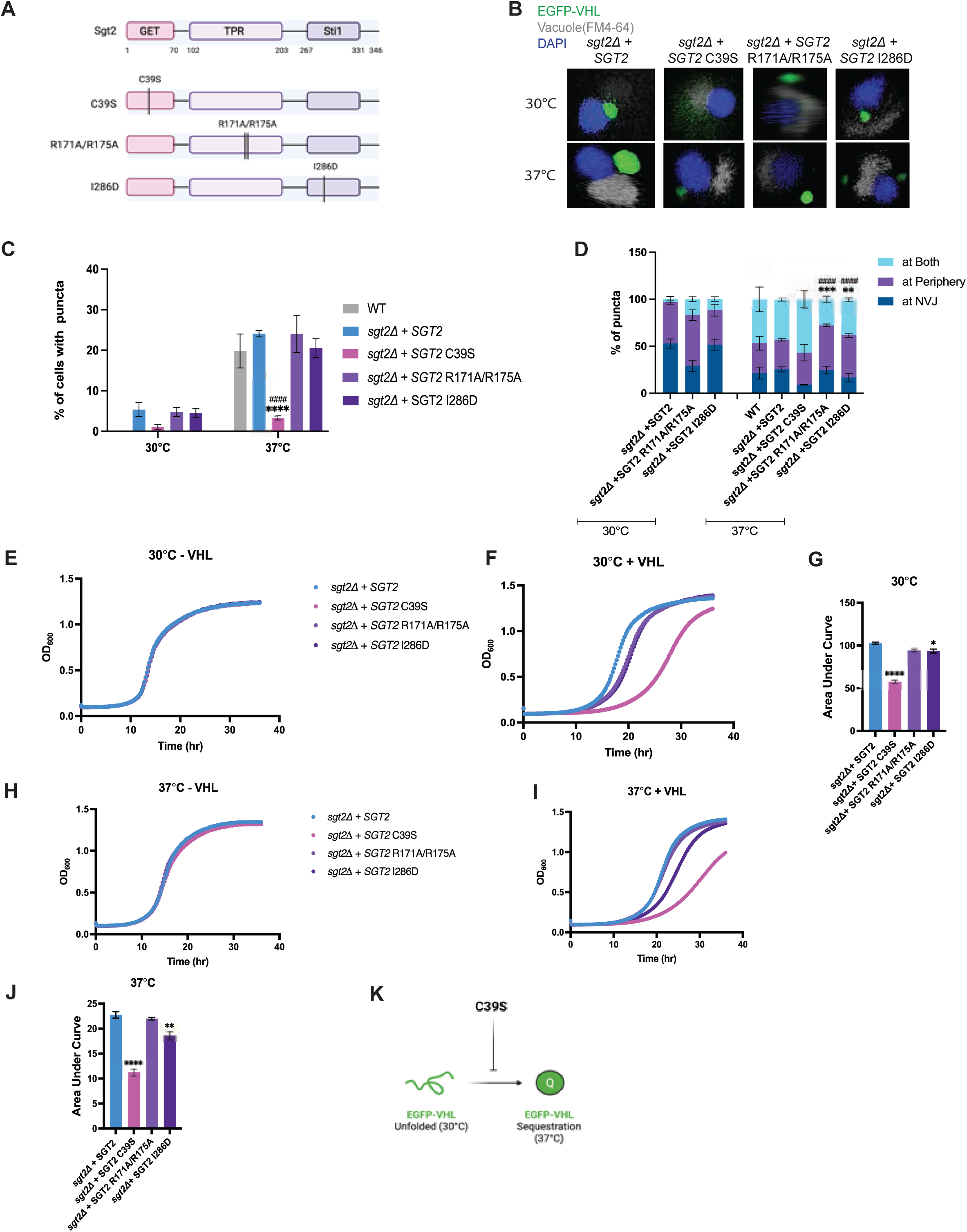
Blocking GET protein binding to Sgt2 inhibits Q-body formation. (A) schematic showing the domains of Sgt2 protein and the point mutations created to block protein interactions with each domain. (B) Representative images of FM4-64 vacuole staining of live cells (gray) with Hoechst 3342 staining (blue) prior to and after 30 minutes of heat shock. (C) Percentage of cells with puncta over time. Fisher’s test: ****p-value<0.0001. Statistics were not run at 30°C due to low percentage of cells with inclusions. (D) Quantification of inclusion localization from (C). Inclusions in proximity to the nuclear-vacuolar junction were categorized at NVJ. Inclusions away from nuclear-vacuolar junction on either the nuclear or vacuolar side were categorized as at periphery. Cells with inclusions at least one inclusion near the nuclear-vacuolar junction and at least one inclusion away from the nuclear-vacuolar junction were categorized as both. Fisher’s test and Bonferroni correction: *sgt2Δ* + *SGT2* vs. *sgt2Δ* + *SGT2* R171A/R175A ***p-value=0.0002, *sgt2Δ* + *SGT2* vs. *sgt2Δ* + *SGT2* I286D **p-value=0.0016, WT vs *sgt2Δ* + *SGT2* R171A/R175A ####p-value<0.0001, WT vs *sgt2Δ* + *SGT2* I286D ####p-value<0.0001. Yeast growth of wildtype *SGT2* mutant strains in 2% galactose and 2% raffinose media at 30°C without VHL present (E), at 30°C with VHL present (F), at 37°C without VHL present (H), and 37°C with VHL present (I). Area under the curve analysis of (F) in (G) and (I) in (HJ. In (G), one-way ANOVA: *sgt2Δ* + *SGT2* vs. *sgt2Δ* + *SGT2* C39S ****p-value<0.0001, *sgt2Δ* + *SGT2* vs. *sgt2Δ* + *SGT2* R171A/R175A *p-value=0.0389, *sgt2Δ* + *SGT2* vs. *sgt2Δ* + *SGT2* I286D *p-value=0.0239. In (J), one-way ANOVA: *sgt2Δ* + *SGT2* vs. C39S ****p-value<0.0001, *sgt2Δ* + *SGT2* vs. *sgt2Δ* + *SGT2* I286D **p-value=0.0043. (I) schematic showing C39S mutant inhibits EGFP-VHL inclusion formation.

### Blocking the GET binding domain of Sgt2 reduces TA protein inclusion formation

Because we saw an increase in toxicity and a reduction in VHL inclusion formation in the Sgt2 C39S GET domain mutant, we hypothesized the mutant could be blocking Sed5 inclusion formation. This could be detrimental to the cell because Sed5 has a role in PQC compartment trafficking leading to compartment degradation^36^. We co-expressed GFP-Sed5 with the Sgt2 C39S mutant. While we did see an increase in diffuse Sed5 in the C39S mutant indicative of Sed5 mislocalization, it was not dramatically different from *sgt2Δ* or *SGT2* overexpression cells (Fig. 6b-d).

**Figure 6:**
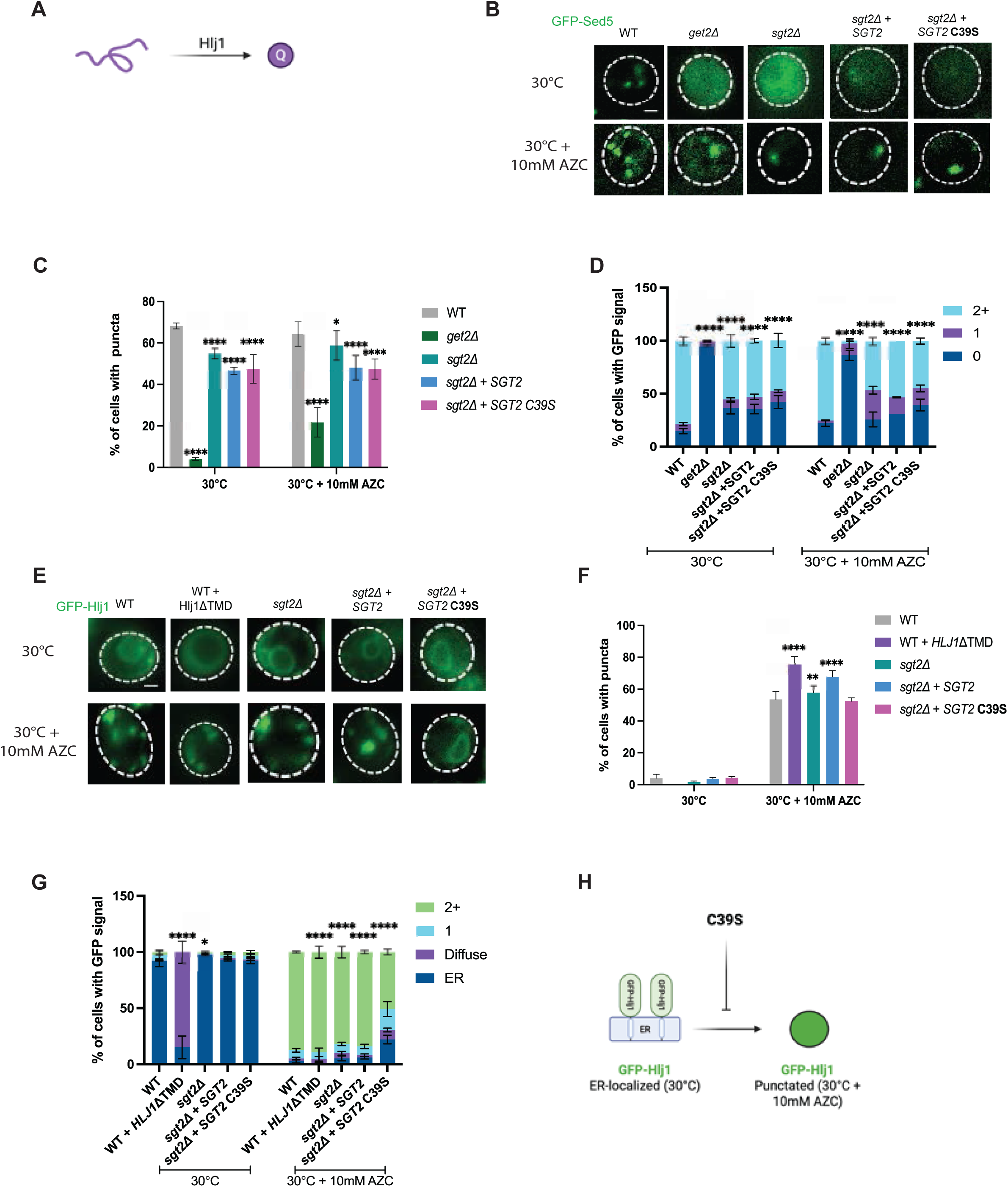
Blocking GET protein binding to Sgt2 reduces inclusion formation of Q-body chaperones. (A) schematic showing the tail anchored J-domain protein Hlj1 aids in Q-body formation. (B) Live cell microscopy of wildtype, *GET2* deletion, and *SGT2* deletion, *SGT2* overexpression, and *SGT2* C39S mutant strains expressing GFP-Sed5 +/- 1 hour of 10mM AZC treatment. Representative images are shown. Three biological replicates were performed. (C) Percentage of cells with puncta over time. Fisher’s test and Bonferroni correction: ****p-value<0.0001. (D) Categorization of number of inclusions in cells expressing GFP-Sed5. Fisher’s test and Bonferroni correction: ****p-value<0.0001. (E) Live cell microscopy of wildtype, *GET2* deletion, and *SGT2* deletion, *SGT2* overexpression, and *SGT2* C39S mutant strains expressing GFP-Hlj1 +/- 1 hour of 10mM AZC treatment. Representative images are shown. Three biological replicates were performed. (F) Percentage of cells with puncta over time. Fisher’s test and Bonferroni correction: WT + AZC vs. WT + Hlj1ΔTMD + AZC ****p-value<0.0001, WT + AZC vs. *sgt2Δ* + AZC **p-value=0.0044, WT + AZC vs. *sgt2Δ*+*SGT2*+AZC ****p-value<0.0001. Statistics were not run on WT + Hlj1ΔTMD without AZC treatment due to lack of variance. (G) Categorization of number of inclusions in cells expressing GFP-Sed5. Fisher’s test and Bonferroni correction: WT vs. WT + Hlj1ΔTMD ****p-value<0.0001, WT vs. *sgt2Δ* *p-value=0.0444, WT + AZC vs. WT + Hlj1ΔTMD + AZC ****p-value<0.0001, WT + AZC vs. *sgt2Δ* + AZC ****p-value<0.0001, WT + AZC vs. *sgt2Δ* + *SGT2* + AZC ****p-value<0.0001, WT + AZC vs. *sgt2Δ* + *SGT2* C39S ****p-value<0.0001. (H) schematic showing C39S mutant prevents Hlj1 inclusion formation during proteotoxic stress.

We next wondered if the C39S mutant influenced chaperones important for inclusion formation, such as J-domain proteins. We chose to investigate Hlj1 because it is a J-domain protein necessary for Q-body formation^4^ (Fig. 6a) and is also a tail anchored protein^9^ localized to the ER (Fig. S6a). We expressed GFP-Hlj1 in wild type and *sgt2Δ* cells and co-expressed GFP-Hlj1 with the *SGT2* overexpression plasmid and its C39S counterpart. As a negative control, we also expressed GFP-Hlj1 lacking its transmembrane domain (TMD) in wild type cells. The GFP-Hlj1ΔTMD construct is known for being diffuse in the cytosol.

We did not observe differences in Hlj1 localization under unstressed conditions (Fig. 6e-g) or major changes in Hlj1 steady state levels (Fig. S6b). When we treated wild type cells with 10mM AZC, we noticed GFP-Hlj1 formed inclusions around the cell periphery like Q-bodies. When we treated with AZC in *sgt2Δ* cells, *SGT2* overexpression cells, and cells expressing the C39S mutant, we observed significant changes in GFP-Hlj1 localization from wild type cells. Most dramatically, the percent of cells with GFP signal with multiple Hlj1 puncta decreases from 95% in wild type cells to less than 50% in cells expressing the C39S mutant. While a large proportion of cells form just one punctum like in Sed5 cells, we noticed that in over 20% of cells Hlj1 is still localized to the ER but does not form puncta in the C39S mutant. Interestingly, when we delete the transmembrane domain of Hlj1, Hlj1 still forms puncta under AZC treatment (Fig. 6e-g) suggesting the C39S mutant may be blocking Q-body formation through Hlj1 in a manner independent of Hlj1 localization (Fig. 6h).

### Other GET pathway chaperones alter VHL inclusion formation

Because reducing Sgt2’s interaction with GET pathway chaperones in the C39S mutant greatly reduced inclusion formation of both GFP-VHL and GFP-Hlj1, and we observed a single Sed5 punctum in a subset of *get2Δ* cells, we wanted to determine if other GET pathway chaperones could alter VHL inclusion formation. In the same manner, we expressed EGFP-VHL into wild type, *get3Δ*, and *get2Δ* cells and tracked GFP-VHL inclusion formation before and after heat shock at 37°C. We chose *GET3* and *GET2* because they are both utilized at different steps in the GET pathway from *SGT2* and each other in the tail anchored protein insertion pathway (Fig. 7a). *get3Δ* and *get2Δ* cells formed GFP-VHL inclusions prior to heat shock to comparable levels as *sgt2Δ* cells (Fig. 7b-d). Cells primarily formed 1-2 large inclusions even more drastically than in *sgt2Δ* cells (Fig. 7d). Interestingly, we observed a significant decrease in the formation of GFP-VHL inclusions after heat shock in *get2Δ* cells compared to wild type cells (Fig. 7b-c) like our C39S mutant. Loss of *GET3* or *GET2* reduces growth, but this reduction appears to be dependent on heat shock not on the presence of VHL (Fig. 7e-h, S7b-c). This suggests that an inefficient GET pathway at any stage of tail anchored protein insertion increases VHL sequestration (Fig. 7i).

**Figure 7:**
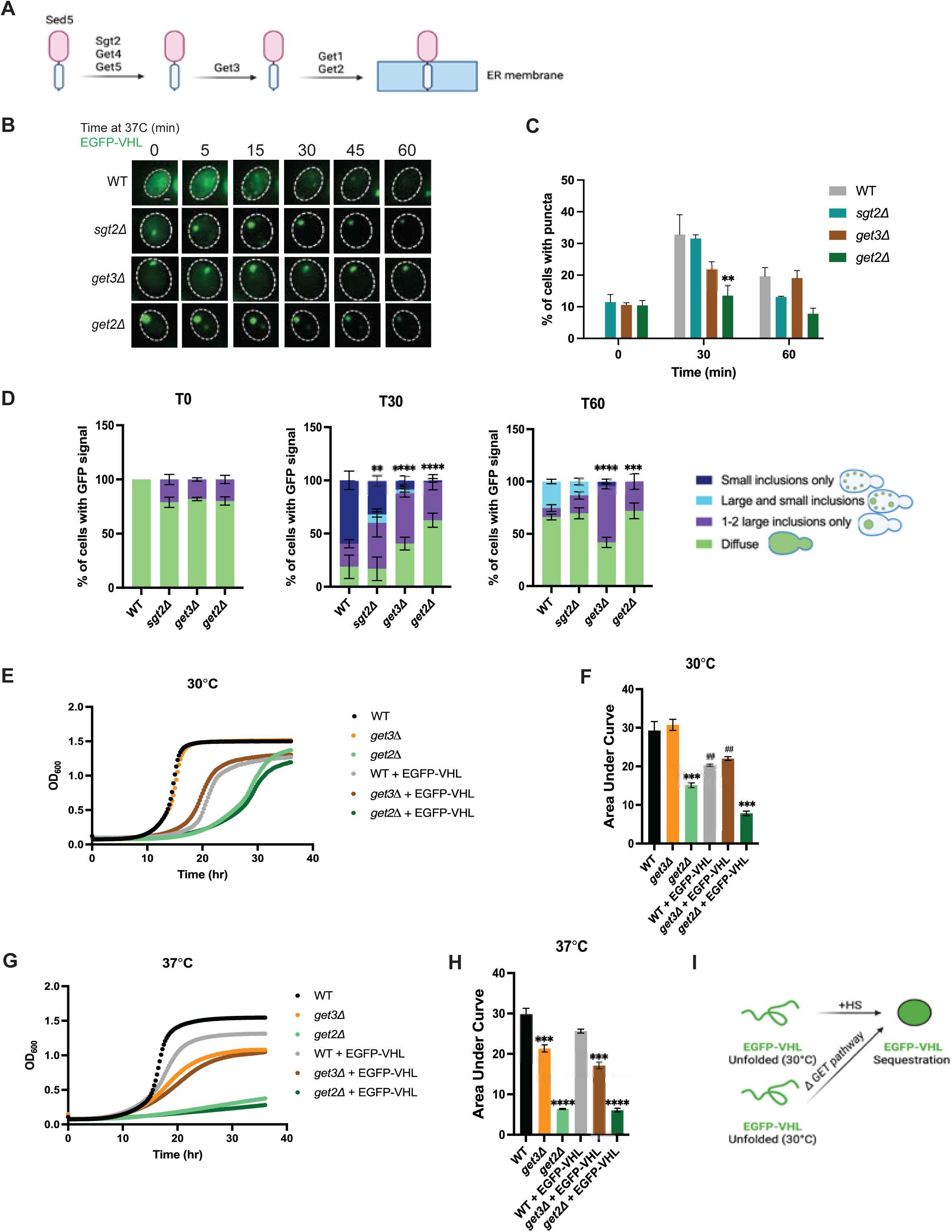
Loss of *GET2* reduces Q-body formation. (A) schematic showing the GET pathway chaperones involved in tail anchored protein insertion at the ER. (B) Time-resolved live cell microscopy of wildtype, *GET3* deletion, *GET2* deletion, and *SGT2* deletion strains expressing EGFP-VHL at 37°C (green). Representative still frames at the times shown. Three biological replicates were performed. (C) Percentage of cells with puncta over time. Fisher’s test and Bonferroni correction: at T30 **p-value=0.0095. (D) Categorization of inclusions in cells expressing EGFP-VHL at 37°C after 30 minutes and 60 minutes of timelapse. Fisher’s test and Bonferroni correction: at T30 WT vs *sgt2Δ* **p-value=0.005, WT vs. *get3Δ* ****p-value<0.0001, WT vs. *get2Δ* ****p-value<0.0001. At T60 WT vs *get3Δ* ****p-value<0.0001, WT vs. *get2Δ* ***p-value=0.0002. Yeast growth of wildtype BY4741 and *SGT2* deletion strains in 2% galactose and 2% raffinose media at 30°C (E) and 37°C (G). Area under the curve analysis of (E) in (F) and (G) in (H). In (F), WT vs. *get2Δ* ***p-value=0.003, WT vs. WT + EGFP-VHL ##p-value=0.0051, *get3Δ* vs. *get3Δ* + EGFP-VHL ##p-value=0.0062, WT + EGFP-VHL vs. *get2Δ* + EGFP-VHL ***p-value=0.001. (I) schematic showing loss of the GET pathway can induce premature inclusion formation of EGFP-VHL.

Work from Kohl and colleagues^31^ has demonstrated the overexpression of *SGT2* in *get3Δ* cells reduces growth compared to wild type yeast by overwhelming the GET pathway (Fig. 8a). We therefore wondered if clogging the GET pathway in this way could reduce VHL inclusion formation comparable to what we observed in our C39S mutant and in *get2Δ* cells. When we co-expressed EGFP-VHL with the *SGT2* overexpression plasmid in *get3Δ* cells, we observed a reduction in inclusion formation (Fig. 8b-d). Inclusions that did form predominantly formed 1-2 large inclusions (Fig. 8d). In line with Kohl and colleagues, we found overexpression of *SGT2* in *get3Δ* cells reduced growth. We also found that VHL presence at 37°C increases this toxicity (Fig. 8e-h, S7d-e). Together this data suggests manipulating the GET pathway has direct implications on the formation of protein quality control compartments (Fig. 8i).

**Figure 8:**
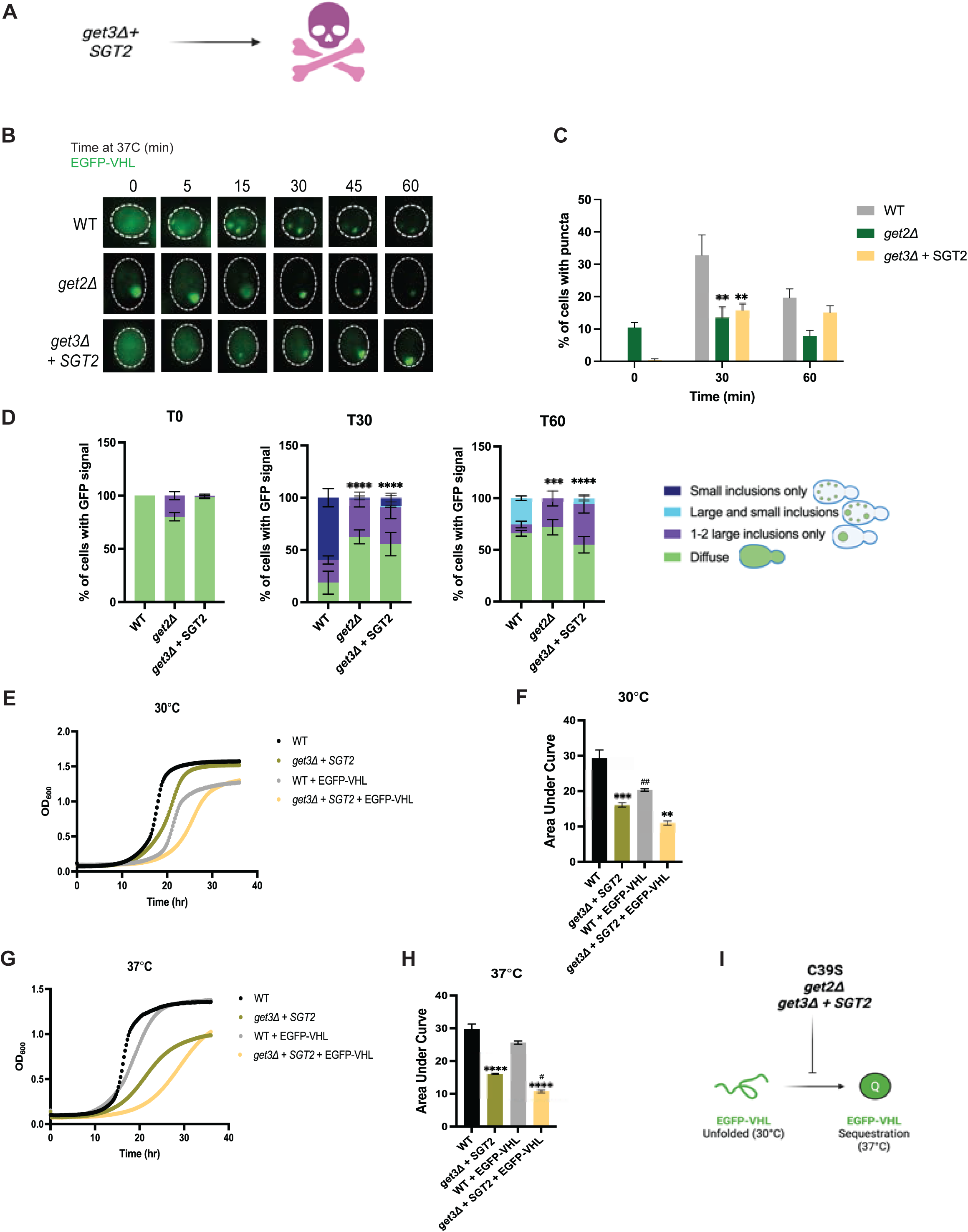
Overexpression of *SGT2* coupled with loss of *GET3* reduces Q-body formation. (A) schematic showing overexpression of *SGT2* in *get3Δ* cells leads to defects in cell growth. (B) Time-resolved live cell microscopy of wildtype, *GET2* deletion, and *GET3* deletion with *SGT2* overexpression strains expressing EGFP-VHL at 37°C (green). Representative still frames at the times shown. Three biological replicates were performed. (C) Percentage of cells with puncta over time. Fisher’s test and Bonferroni correction: at T30 *get2Δ* **p-value=0.0095 *get3Δ*+*SGT2* *p-value=0.0105. (D) Categorization of inclusions in cells expressing EGFP-VHL at 37°C after 30 minutes and 60 minutes of timelapse. Fisher’s test and Bonferroni correction: at T30 *get2Δ* ****p-value<0.0001 and *get3Δ*+*SGT2* ****p-value<0.0001, at T60 *get2Δ* ***p-value=0.0002 and *get3Δ*+*SGT2* ****p-value<0.0001. Yeast growth of wildtype BY4741 and *SGT2* deletion strains in 2% galactose and 2% raffinose media at 30°C (E) and 37°C (G). Area under the curve analysis of (E) in (F) and (G) in (H). In (F), WT vs *get3Δ*+*SGT2* ***p-value=0.0003, WT vs WT + EGFP-VHL ##p-value=0.004. In (H), WT vs *get3Δ*+*SGT2* ****p-value<0.0001, WT+EGFP-VHL vs *get3Δ*+*SGT2*+EGFP-VHL ****p-value<0.0001, *get3Δ*+*SGT2* vs *get3Δ*+*SGT2*+EGFP-VHL #p-value=0.0119. (I) schematic showing EGFP-VHL sequestration can be reduced by C39S mutant, *get2Δ*, or *get3Δ*+*SGT2*.

## Discussion

Here we have identified a role for Sgt2 and other members of the GET pathway in PQC compartment formation through tail anchored protein insertion and have demonstrated for the first time that spatial quality control is required for cell health (Fig. 9). Altering Sgt2 or Get2 levels in any capacity increases the mislocalization of GET pathway substrates to the cytosol promoting their aggregation under proteotoxic stress conditions. These aggregated substrates can promote compartment formation that would not normally occur. It will be interesting to dissect the specificity of this process. Perhaps ER destined tail anchored proteins are ideal candidates for this process because of their more hydrophobic tails in comparison to mitochondrial tail anchored proteins. It is also possible that Sed5 seeds this process due to its role in IPOD trafficking^36^. In either case, it appears to be dependent on an inefficient GET pathway being unable to properly insert Sed5.

**Figure 9:**
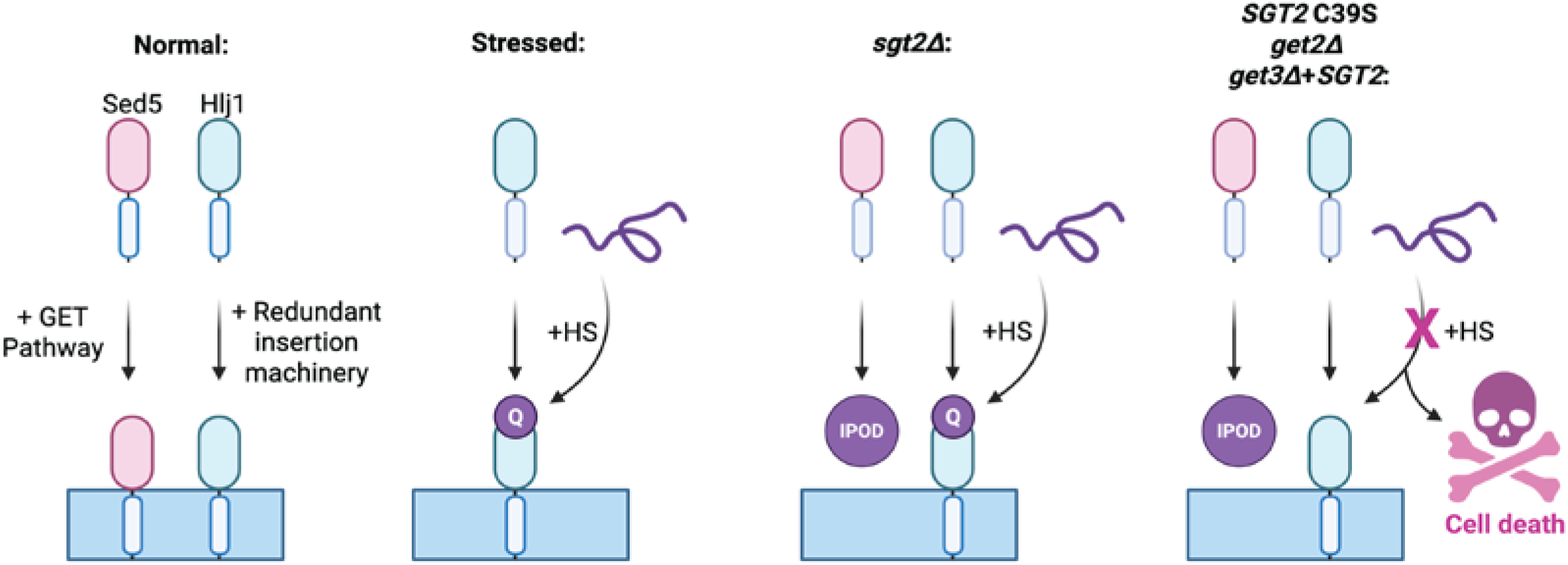
A functional GET pathway is necessary and required for spatial quality control in the cytoplasm. Under wildtype conditions, tail anchored proteins are inserted into the ER membrane by the GET pathway and redundant insertion machinery. Under proteotoxic stress, the insertion of J-domain proteins like Hlj1 allows for the sequestration of misfolded protein into Q-bodies. When the GET pathway becomes less efficient (*sgt2Δ*) mislocalized tail anchored proteins can co-aggregate with misfolded proteins in the IPOD, but Q-bodies still form. When the GET pathway becomes defective (C39S, *get2Δ, get3Δ+SGT2*) Q-bodies do not form leading to cell toxicity.

When we transition the GET pathway from an inefficient to defective state, the PQC pathway no longer functions as it should. Q-bodies no longer form, contributing to growth defects only during proteotoxic stress. In Sgt2 C39S GET binding domain mutant cells, we observed a dramatic reduction in inclusions of the Q-body forming chaperone Hlj1. While we do see increases in mislocalized Hlj1, a large proportion of Hlj1 is at the ER but is not forming inclusions as we expected. Because Hsp70s are required for Q-body formation^2,4^, and Hsp70s deliver tail anchored proteins to Sgt2^19,37^, future studies will determine if Sgt2 regulates Hsp70s by the GET pathway. It is possible that the C39S mutant inhibits Hlj1 inclusion formation by sequestering Hsp70s. If this is the case, our data effectively demonstrates the consequence of the same major chaperones being involved in so many processes like post-translational targeting, protein folding, and sequestration. This also highlights how necessary it is to regulate major chaperone function.

The reductions in VHL inclusion formation were replicated by *get2Δ* cells and *SGT2* overexpression in *get3Δ* cells. While we saw decreases in inclusion formation that mirrored our C39S mutant, we did see a larger proportion of cells that formed inclusions but only formed 1-2 per cell. It is possible that because our C39S mutant also had decreased VHL expression, we would see a similar 1-2 inclusions with higher VHL expression to the levels seen in the *get2Δ* and *SGT2* overexpression in *get3Δ* cells. In each case, we see a reduction in Q-body formation when we break the GET pathway at any stage. With the reductions in Q-body formation, we observe cellular toxicity. It should be noted that this toxicity does not exist unless VHL is present suggesting that cellular toxicity is an effect of the change in spatial quality control not of the *get* deletion alone or simply a result of heat shock.

Our findings connect protein biogenesis and spatial PQC. While several studies have identified members of the GET pathway as important for PQC^22–24,27^, we have identified a role for tail anchored protein insertion in the function of spatial PQC compartments. Our findings raise the possibility that errors in protein insertion can impair the sequestration of misfolded proteins into PQC compartments. A better understanding of the GET pathway could be necessary for the development of therapies for mitigating these errors as they increase in aging and disease.

More broadly, our findings show that spatial quality control is necessary and required for surviving proteotoxic stress in yeast cells. The JUNQ and IPOD compartments were discovered almost two decades ago5 with the discovery of Q-bodies soon following4. Several researchers have pushed the knowledge of these compartments forward1,2,36,38–41; however, the biological relevance of these compartments has remained unclear. We have shown for the first time that the cell cannot endure proteotoxic stress unless these compartments form. It is well established that the IPOD sequesters amyloids associated with neurodegenerative diseases like expanded exon 1 huntingtin3,5,6. Our work highlights the importance and urgency of targeting PQC compartments in protein misfolding diseases. It is very possible that the toxicity associated with neurodegenerative diseases is contributed by the dysregulation of PQC compartments.

## Methods

### Plasmids and yeast strains

Plasmids used in this study are summarized in Table 1. NES-GFP-LuciferaseTs (NES-LuciTs) was expressed using a galactose promoter in pAG426^2,3^. EGFP-VHL was expressed using a galactose promoter in pESC^1^. GFP-Sed5 was expressed using an ADH1 promoter in pRS315 and was a gift from Blanche Schwappach. mCherry-Sed5 was expressed using a MET25 promoter in pRS415 and was a gift from Blanche Schwappach. GFP-Hlj1 was expressed using a MET25 promoter in pRS416 and was a gift from Jeffery Brodsky. Sgt2-mCherry and Sgt2-mCherry point mutants were expressed using a galactose promoter in pAG413. Sgt2-mCherry was constructed from pDONR Sgt2 purchased from Horizon (cat. YSC3867-202329690) and pAG413GAL-ccdB-DsRed was a gift from Susan Lindquist (Addgene plasmid # 14357). Point mutants were constructed using site-directed mutagenesis.

**Table 1:**
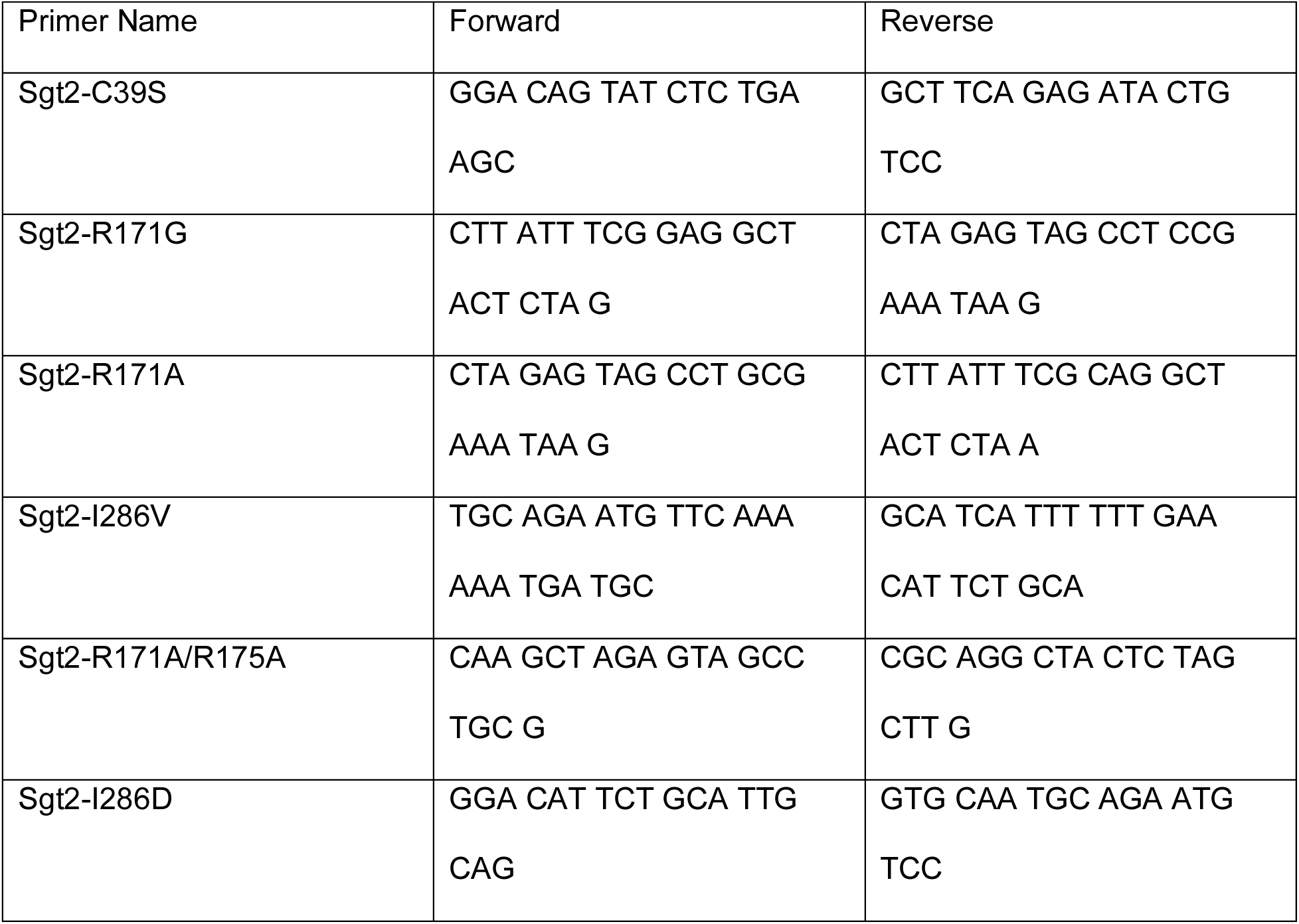
List of primers used.

**Table 2:** List of plasmids used.

| Plasmid Name | Backbone | CEN or 2u | From |
| --- | --- | --- | --- |
| NES-GFP-LuciferaseTs | pAG426 | 2u | <sup>1,2</sup> |
| EGFP-VHL | pESC | 2u | <sup>1</sup> |
| GFP-Sed5 | pRS315 | CEN | Gift from Blanche Schwappach |
| GFP-Hlj1 | p416MET25 | CEN | Gift from Jeffrey Brodsky |
| GFP-Hlj1ΔTMD | p416MET25 | CEN | Gift from Jeffrey Brodsky |
| pENTR-Sgt2 | pENTR | pENTR | Horizon (cat. YSC3867-202329690) |
| pENTR-Sgt2 C39S | pENTR | pENTR | This paper |
| pENTR-Sgt2 R171A/R175A | pENTR | pENTR | This paper |
| pENTR-Sgt2 I286D | pENTR | pENTR | This paper |
| Gal-ccdB-CHFP-HIS | pAG413 | CEN | Gift from Susan Lindquist |
| Sgt2-mCherry | pAG413 | CEN | This paper |
| Sgt2-mCherry C39S | pAG413 | CEN | This paper |
| Sgt2-mCherry R171A/R175A | pAG413 | CEN | This paper |
| Sgt2-mCherry I286D | pAG413 | CEN | This paper |

### Time-resolved live cell imaging

Yeast cultures were grown as previously described^1,2^. Briefly, yeast was grown at 30°C to stationary phase in selective media containing 2% raffinose and then diluted to OD600 0.1 in selective media containing 2% raffinose and 2% galactose. Cultures grew at 30°C for 4 hours to induce expression of GAL-inducible proteins. Cells were adhered to coverslips containing concanavalin A (Sigma-Alrich cat #C2010). Cells were washed with selective media containing 2% glucose and kept in the same medium by sealing the coverslips with vacuum grease. Cells were imaged every 15s or 1min as indicated for 1 hour at 37°C using Zeiss Axio Observer.Z1 7 inverted microscope (RRID:SCR_023694) equipped with X-Cite Xylis LED light source (EXCELITA Technologies), HE GFP/Cy3/DAPI shift free filter sets (Zeiss), a Plan-Apochromat 100x/1.40 oil DIC M27 objective (Zeiss), and a digital Axiocam 705 camera (Zeiss) controlled by Zen blue software (RRID:SCR_013672).

### FM4-64 vacuole labeling

Cells were grown as described for time-resolved imaging. After 3.5 h of growth in selective galactose media, cells were resuspended in fresh selective galactose media. FM4-64 (Invitrogen Cat #T3166) was added to a final concentration of 8 μM, and cells were incubated at 30°C for 30 min. Cells were pelleted and resuspended in selective media containing 2% glucose without FM4-64. Cells were incubated for 1 hour at 30°C or 37°C as indicated. Hoescht 33342 (Invitrogen Cat# H3570) was added prior to imaging to a final concentration of 8.1 μM to visualize nucleic acid. Cells were adhered to coverslips using concanavalin A (Sigma-Aldrich Cat #C2010-250MG) and washed with 100 mM HEPES pH 7.4 with 5% glucose. Images were taken using z-stacks, taking 10 stacks with a step size of 1 μm. Images were taken using Zeiss Axio Observer.Z1 7 inverted microscope described above Inclusions in proximity to the nuclear-vacuolar junction were categorized at NVJ. Inclusions away from nuclear-vacuolar junction on either the nuclear or vacuolar side were categorized as at periphery. Cells with inclusions at least one inclusion near the nuclear-vacuolar junction and at least one inclusion away from the nuclear-vacuolar junction were categorized as both.

### Growth curve assays

Yeast cultures were grown at 30°C in selective media containing 2% raffinose to stationary phase and diluted to OD600 0.1 in selective media containing 2% raffinose. Cultures were further diluted to a final OD600 0.01 in selective media containing 2% raffinose and 2% galactose or 2% glucose in a round bottom 96 well plate (Thermo Fisher cat #650 185). Cultures grew for 36 hours with constant shaking. OD600 measurements were taken every 15 min using Molecular Devices SpectraMax Mini Microplate Reader. Area under the curve was calculated using GraphPad Prism 10.0.1 (RRID:SCR_002798) and compared using a one-way ANOVA with Tukey’s multiple comparisons test.

#### Photobleaching microscopy

Cells were grown as described for time-resolved imaging. After 4 h of growth in galactose media, cells were heat shocked for 1 h at 37°C. Mutant huntingtin exon 1 protein (HTT) was used as a control for IPOD formation and photobleaching. HTT expressing cells were grown in 2% galactose, 2% raffinose for 24 h at 30°C prior to imaging. Cells were adhered to coverslips using concanavalin A (Sigma-Aldrich Cat #C2010-250MG). Photobleaching was performed on a Nikon TE2000 Perfect Focus Ti - E microscope with phase contrast, epi-fluorescence, DIC, and resonant scanning confocal system for 405, 488, 561, and 638 nm lasers (RRID:SCR_023161) controlled by Nikon NIS-Elements software (RRID:SCR_014329). Cells were imaged pre-photobleaching, the cytosol of a cell with an inclusion was bleached using the 488 nm laser for 5 s at 100% power, and cells were then imaged every 2 s for 90 s post-bleaching. Photobleaching image analysis was performed using ImageJ. Integrated density measurements using a drawn ROI were normalized to the pre-bleach image.

### Single timepoint live cell imaging

Cells were grown as described for time-resolved imaging. For plasmids with the MET25 promoter, cells were diluted to OD600 0.1 in selective media lacking methionine and containing 2% raffinose and 2% galactose or containing 2% glucose when galactose expression was not required. Cultures were grown for 30°C for 3 hours to induce expression. Cultures were pelleted and resuspended in fresh selective media with or without 10mM L-Azetidine-2-carboxylic acid (Sigma-Alrich cat A0760. Cultures grew at 30°C for 1 hour. Cells were adhered to coverslips containing concanavalin A (Sigma-Alrich cat C2010). Cells were washed with 2% glucose or 1x PBS. Cells were imaged using Zeiss Axio Observer.Z1 7 inverted microscope (RRID:SCR_023694) described above.

### Microscopy image analysis

Images were analyzed using ImageJ (RRID:SCR_003070) and Volocity (RRID:SCR_002668). No alterations were made to the time-lapse images prior to puncta counting analyses. Background subtraction in Zen using a radius of 100, photobleaching correction (v2.1.0 exponential fit)^29^ and registration (descriptor-based series (2d/3d + t) nearest neighbor, fuse and display with Σ1 = 12.901999, Σ2 = 15.343149, Threshold = 0.012469863)^30^ were performed on all time lapse data after puncta counting analyses using ImageJ for visualization purposes only. Fisher’s tests were performed in Prism (RRID: SCR_002798) to determine statistical significance. Bonferroni correction was performed when analysis required 3 or more groups. 3D reconstructions were performed in Volocity with adjustments to black and white levels to optimize visualization.

### SDS-PAGE with Western blotting

Yeast cultures were grown at 30°C in selective media containing 2% raffinose and 2% galactose or containing 2% glucose when galactose expression was not required for 20 hours. Cultures were diluted to OD600 0.2 in selective media containing 2% raffinose and 2% galactose or containing 2% glucose when galactose expression was not required and grown at 30°C for 4 hours. Plasmids utilizing the MET25 promoter were diluted in selective media lacking methionine containing 2% glucose at this point. After 4 hours cells were pelleted and frozen at - 80°C.

Cells were lysed in urea lysis buffer (8 M urea, 50 mM HEPES pH 7.4, Roche cOmplete protease inhibitor Cat# 11836170001) and vortexed with glass beads at 4°C for 5 minutes. 20% SDS was added to a final concentration of 4% SDS, and lysate was incubated at 65°C for 5 min. Lysate was cleared by centrifugation, and protein concentration of the supernatant was measured with the bicinchoninic acid protein assay kit (Thermo Fisher Cat# 23250).

25μg of total protein was resolved on 12% Tris-glycine gels and transferred to nitrocellulose (BioRad Cat# 1620112). EGFP was detected with mouse anti-GFP monoclonal antibody (Roche Cat# 11814460001, RRID:AB_390913), DsRed and mCherry were detected with Living Colors rabbit anti-DsRed polyclonal antibody (Takara Bio Cat# 632496, RRID:AB_10013483), HSP70 was detected with mouse anti-HSP70 monoclonal antibody (Invitrogen cat #MA3-006) and GAPDH was detected with rabbit anti-GAPDH monoclonal antibody (GeneTex cat #GTX637966). Secondary antibodies used were rabbit Anti-Mouse IgG (H + L), HRP Conjugate (Promega Cat# W4021 (also W402B), RRID:AB_430834) and goat Anti-Rabbit IgG (H + L), HRP Conjugate (Promega Cat# W4011 (also W401B), RRID:AB_430833). All primary antibodies were diluted 1:5000 using 5% BSA, and all secondary antibodies were diluted 1:10000 using 5% BSA. Membranes were imaged using Clarity Western ECL Substrate (Biorad Cat# 170-5,061) on GE Amersham Imager 600. Band intensities were measured using ImageJ (RRID:SCR_003070) and normalized to the GAPDH control.

## Supporting information

This article contains supporting information.

## Acknowledgements

We would like to thank Dr. Judith Frydman, Dr. Blanche Schwappach, Dr. Jeffrey Brodsky, and Dr. Susan Lindquist for plasmids used in this study.

## Author Contributions

C.A.L designed and performed experiments and data analyses and wrote the manuscript.

E.M.S. designed experiments, edited the manuscript, and managed the project.

## Data Availability

All data are contained within the manuscript and supplemental materials.

## Supplemental Figure Legends

**Figure S1:**
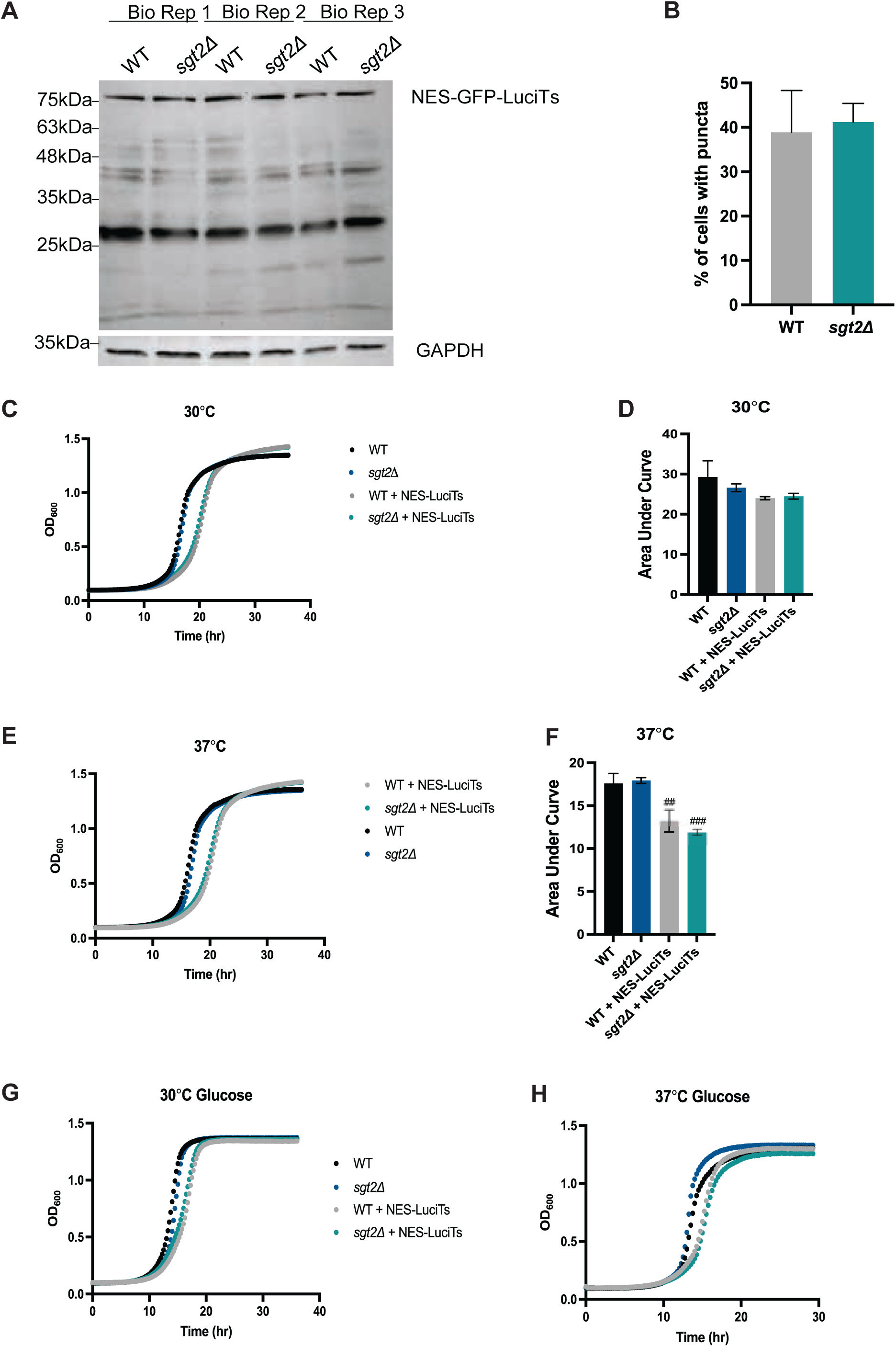
Loss of *SGT2* does not alter NES-LuciTs steady-state protein levels. (A) Western blot comparing steady state levels of NES-LuciTs in wildtype and *sgt2Δ* cells. (B) Percentage of cells with puncta after FM4-64 staining as indicated in Figure 1 to confirm equivalent puncta counts between timelapse and FM4-64 staining.. Fisher’s test: p-value>0.05. Yeast growth of wildtype BY4741 and *SGT2* deletion strains in 2% glucose media at 30°C in (C) and 37°C in (D). Three biological replicates were performed.

**Figure S2:**
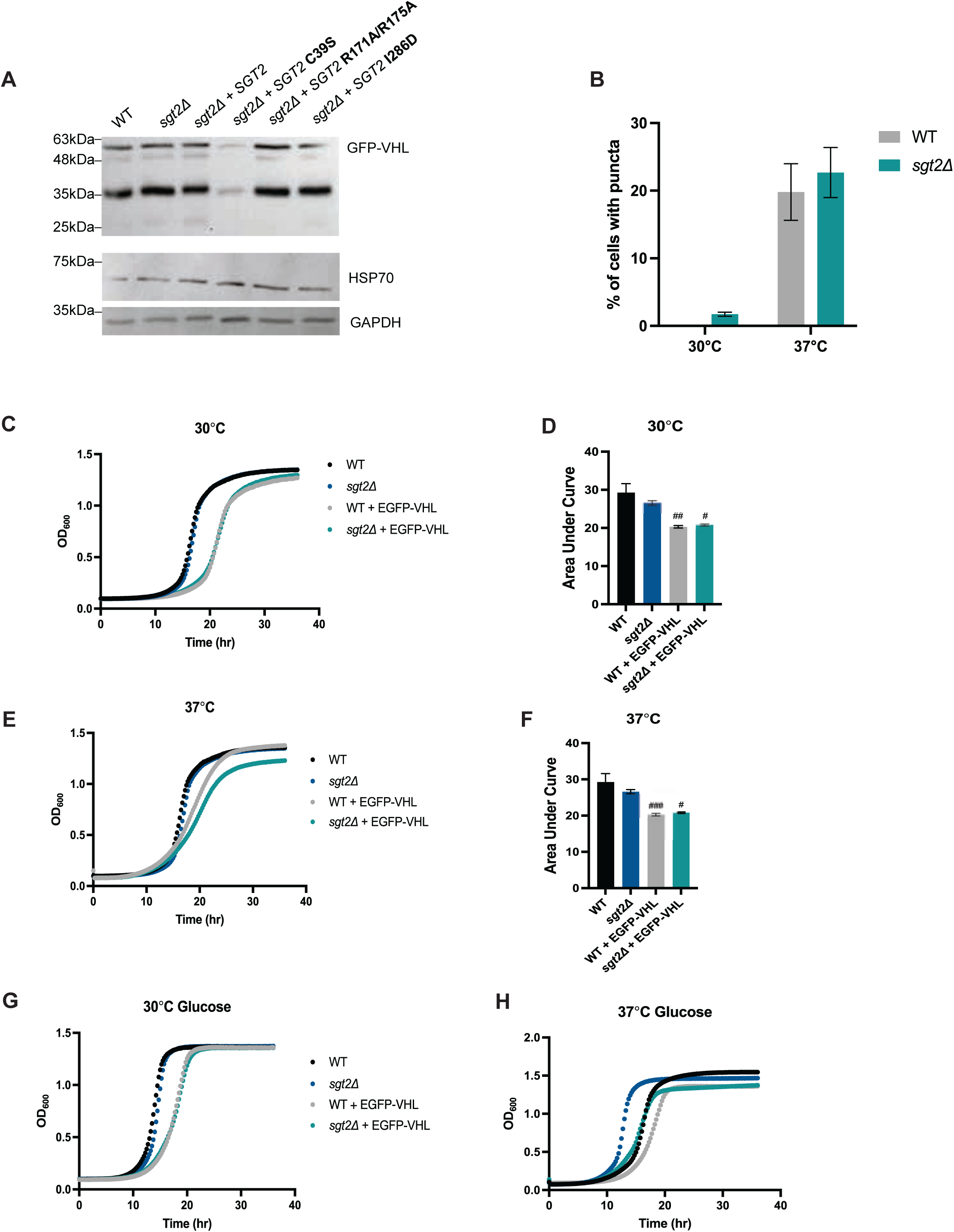
Loss of *SGT2* does not alter EGFP-VHL steady state levels. (A) Western blot comparing steady state protein levels of EGFP-VHL and HSP70 in wildtype, *sgt2Δ*, and *sgt2Δ* cells expressing point mutations. (B) Percentage of cells with puncta after FM4-64 staining as indicated in Figure 2. Fisher’s test: p-value>0.05. Yeast growth of wildtype BY4741 and *SGT2* deletion strains in 2% glucose media at 30°C in (C) and 37°C in (D). Three biological replicates were performed.

**Figure S3:**
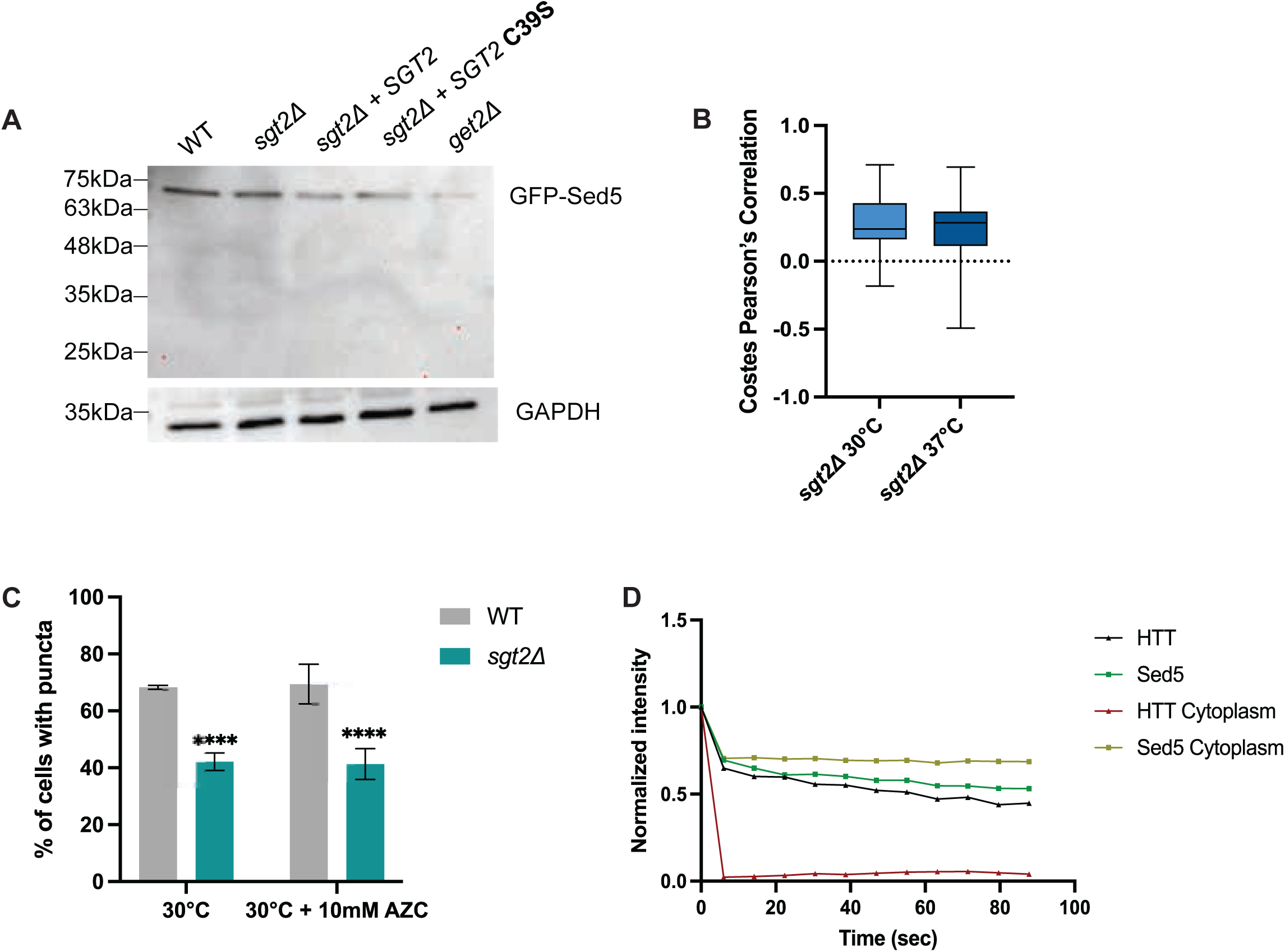
Loss of *SGT2* decreases Sed5 inclusion formation. (A) Western blot comparing steady state levels of GFP-Sed5 in wildtype, *sgt2Δ*, and *sgt2Δ* cells expressing point mutations. (B) Costes Pearson’s correlation co-efficient of mCherry-Sed5 and EGFP-VHL in *sgt2Δ* cells at 30°C and 37°C respectively. (C) Percentage of cells with puncta after FM4-64 staining as indicated in Figure 3. Fisher’s test: ****p-value<0.0001. (D) Normalized intensity of aggregate before and after bleaching adjacent cytosol as indicated in Figure 4. Aggregate intensity was normalized to intensity of aggregate before bleaching. If the fluorescent intensity of background >1, aggregate intensity was also normalized to background. Mean of three biological replicates of at least 10 cells from each replicate was plotted.

**Figure S4:**
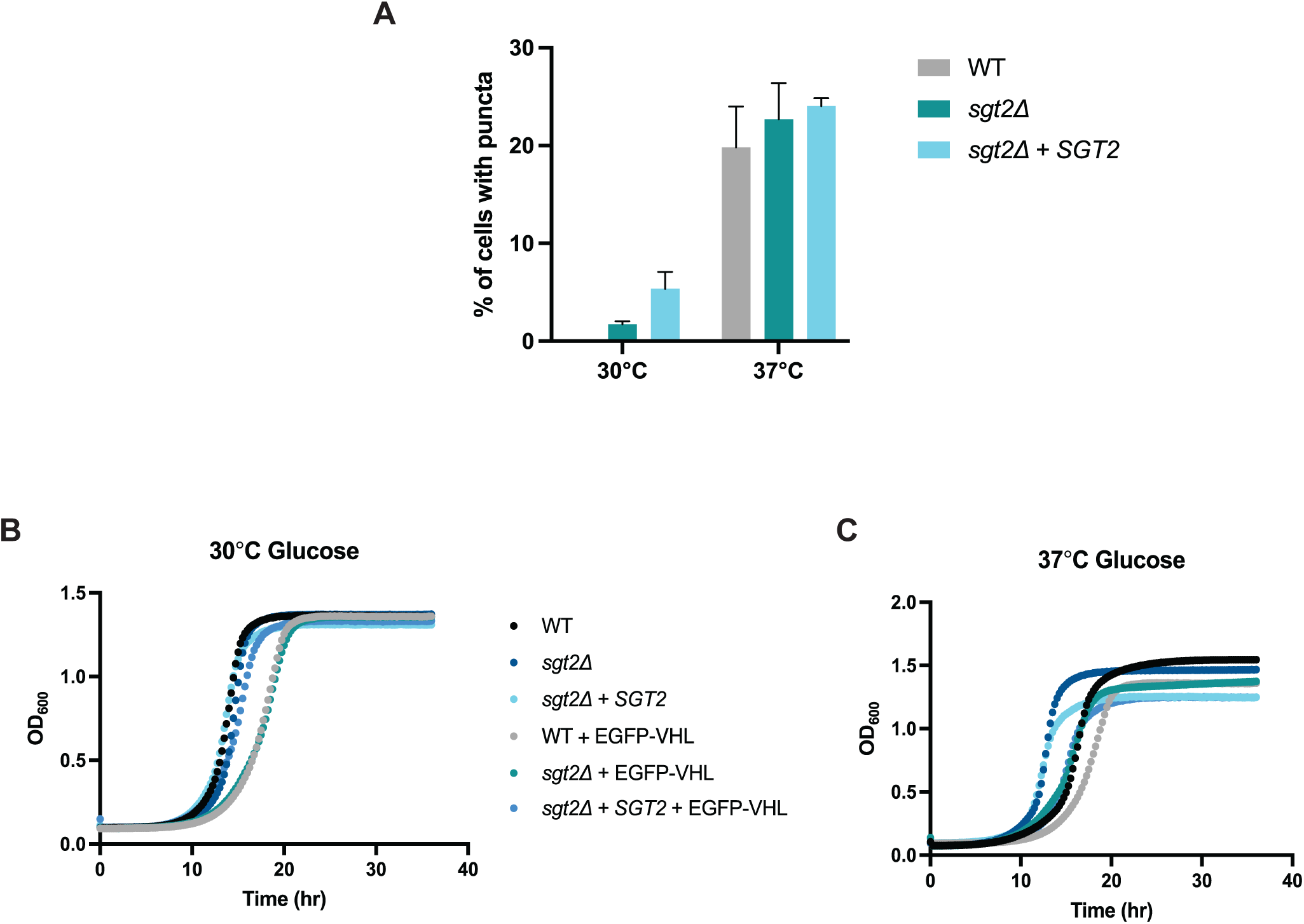
Overexpression of *SGT2* does not significantly alter inclusion formation after heat shock. (A) Percentage of cells with puncta after FM4-64 staining as indicated in Figure 4. Fisher’s test p-value>0.05. Yeast growth of wildtype BY4741, *SGT2* deletion, and *SGT2* overexpression strains in 2% glucose media at 30°C in (B) and 37°C in (C). Three biological replicates were performed.

**Figure S5:**
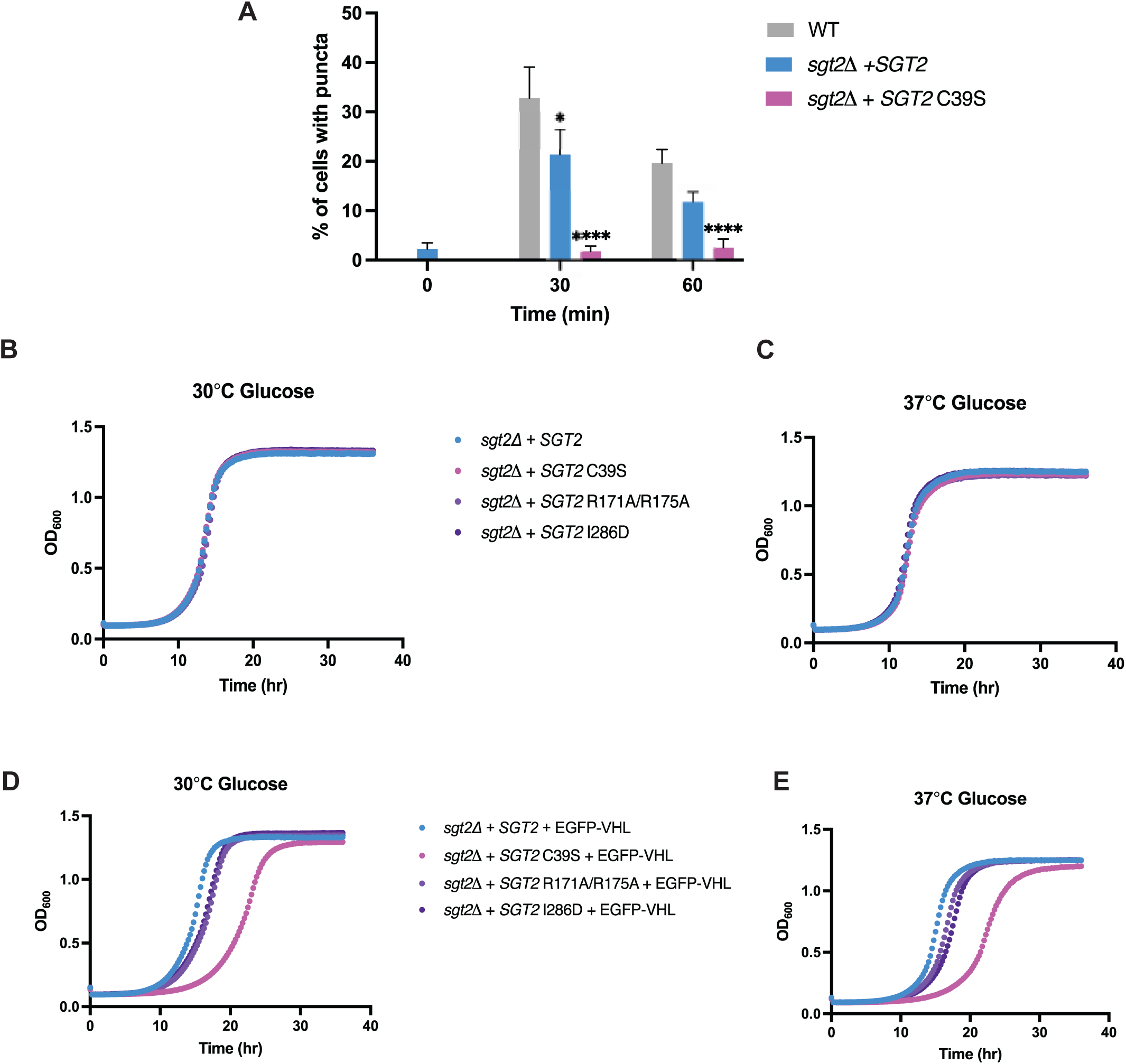
VHL is required to induce toxicity of C39S mutant. Yeast growth of wildtype BY4741, *SGT2* deletion, and *SGT2* overexpression strains in 2% galactose and 2% raffinose media without VHL expression at 30°C in (A) and at 37°C in (B). Yeast growth of wildtype BY4741, *SGT2* deletion, and *SGT2* overexpression strains in 2% glucose media without VHL expression at 30°C in (C) and at 37°C in (D). Yeast growth of wildtype BY4741, *SGT2* deletion, and *SGT2* overexpression strains in 2% glucose media expression at 30°C in (E) and at 37°C in (F). Percentage of cells with puncta during time-resolved live cell microscopy of wildtype BY4741 and *SGT2* overexpression, and C39S mutant strains expressing EGFP-VHL at 37°C. Fisher’s test: ****p-value<0.0001.

**Figure S6:**
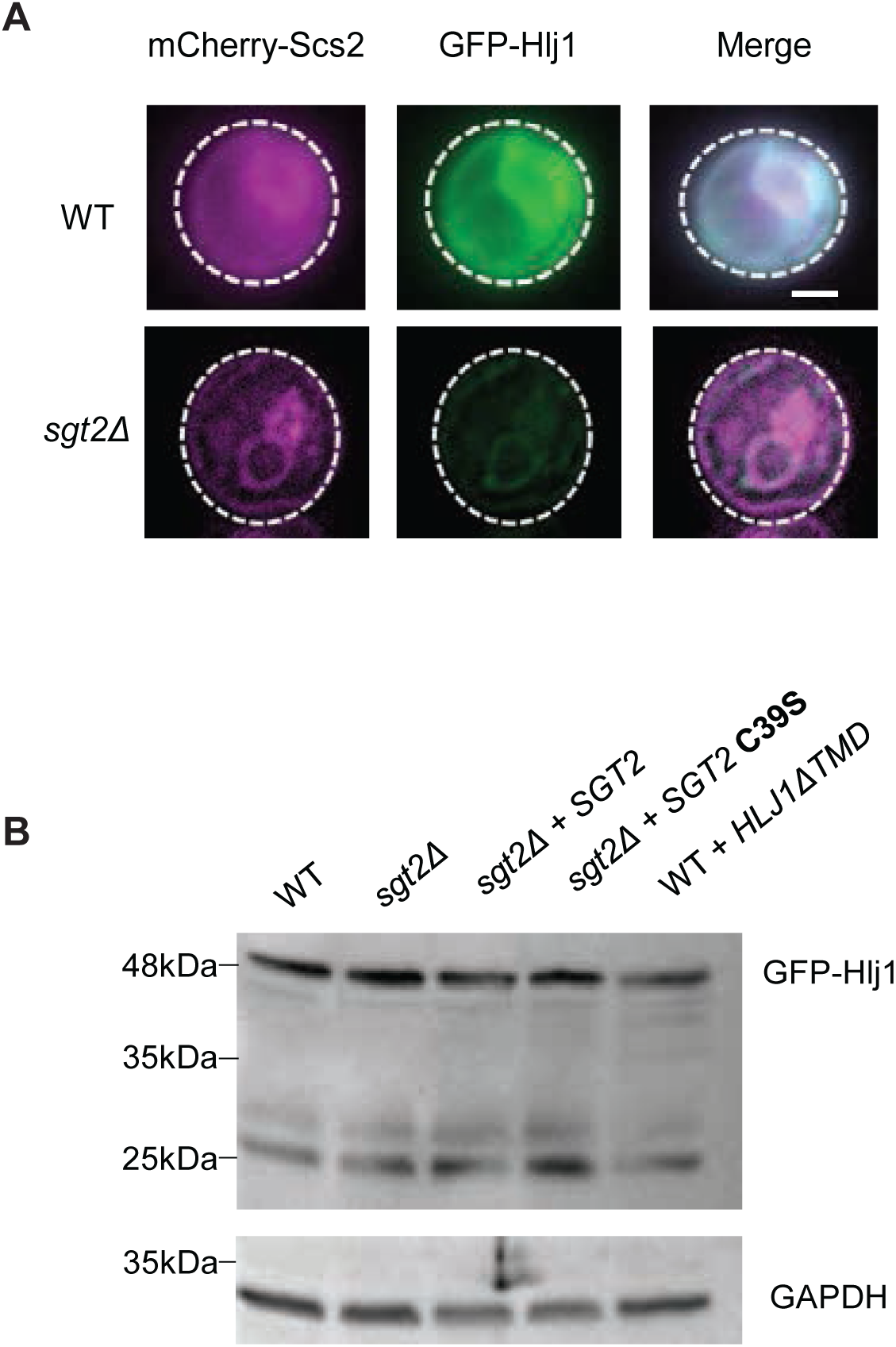
*SGT2* mutants do not alter Hlj1 steady state levels. (A) Colocalization microscopy of ER marker mCherry-Scs2 and GFP-Hlj1 in wildtype and *sgt2Δ* cells. (B) Western blot comparing steady state levels of GFP-Hlj1 in wildtype, *sgt2Δ,* and *sgt2Δ* cells expressing point mutations.

**Figure S7:**
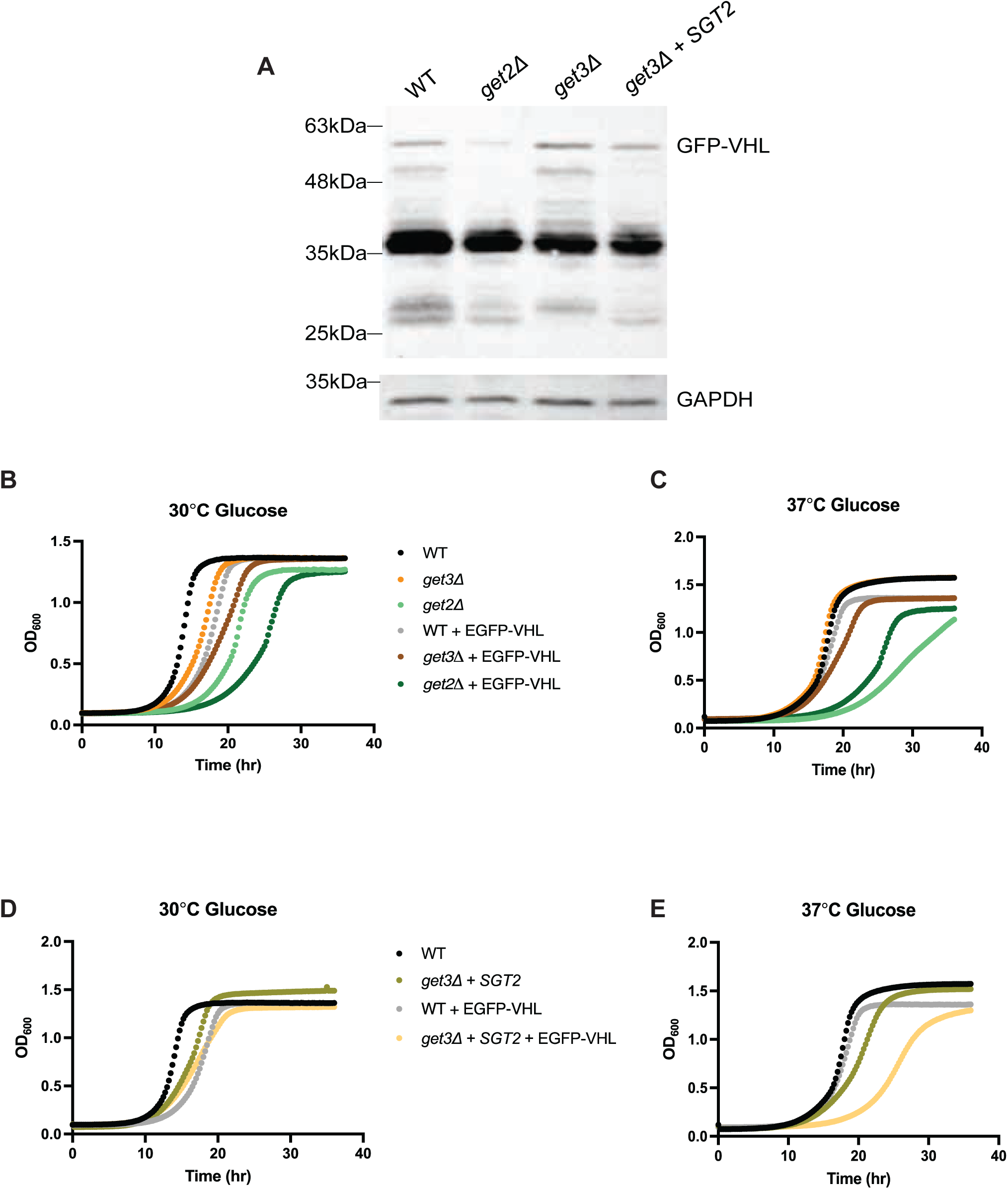
Loss of *GET2* is toxic with or without VHL present. (A) Western blot comparing steady state levels of EGFP-VHL in wildtype, *sgt2Δ, get2Δ, get3Δ,* and *get3Δ+SGT2* strains. Yeast growth of wildtype BY4741, *GET2* deletion, and *GET3* deletion strains in 2% glucose media without VHL expression at 30°C in (B) and at 37°C in (C). Yeast growth of wildtype BY4741 and *get3Δ* coupled with *SGT2* overexpression strains in 2% glucose media without VHL expression at 30°C in (D) and at 37°C in (E).\

**Supplemental Figure 8.**
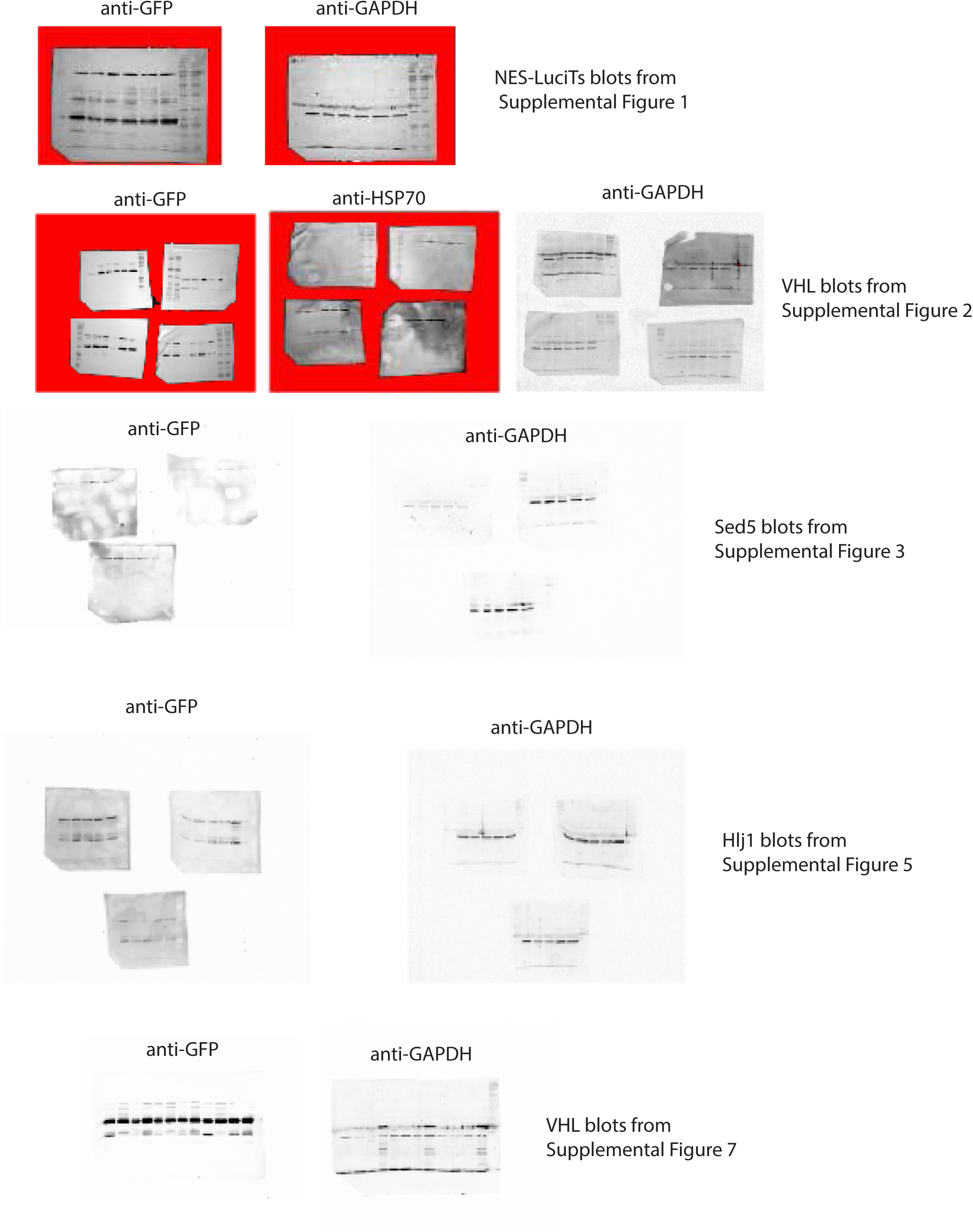

